# Brain network mechanisms of visual shape completion

**DOI:** 10.1101/2020.08.03.233403

**Authors:** Brian P. Keane, Deanna M. Barch, Ravi D. Mill, Steven M. Silverstein, Bart Krekelberg, Michael W. Cole

**Author notes:** Co-senior authors (Bart Krekelberg, Michael W. Cole). Author contributions:* Conceptualization – BPK, DMB, MWC; Data curation – BPK; Formal analysis–BPK, RDM, MWC, DMB; Funding acquisition–BPK; Resources–MWC, BK, SMS; Software–RDM, MWC, BPK; Supervision–MWC; Visualization–BPK, RDM, MWC; First draft – BPK; Review & editing–BPK, RDM, MWC, DMB, BK, SMS. Dedications*: None.

## Abstract

Visual shape completion represents object shape, size, and number from spatially segregated edges. Despite being extensively investigated, the process’s underlying brain regions, networks, and functional connections are still not well understood. To shed light on the topic, we scanned (fMRI) healthy adults during rest and during a task in which they discriminated pac-man configurations that formed or failed to form visually completed shapes (illusory and fragmented condition, respectively). Task activation differences (illusory-fragmented), resting-state functional connectivity, and multivariate pattern differences were identified on the cortical surface using 360 predefined parcels and 12 functional networks composed of such parcels. Brain activity flow mapping (ActFlow) was used to evaluate the likely involvement of resting-state connections for shape completion. We identified 34 differentially-active parcels including a posterior temporal region, PH, whose activity was consistent across all 20 observers. Significant task regions primarily occupied the secondary visual network but also incorporated the frontoparietal, dorsal attention, default mode, and cingulo-opercular networks. Each parcel’s task activation difference could be modeled via its resting-state connections with the remaining parcels (*r*=.62, *p*<10^-9^), suggesting that such connections undergird shape completion. Functional connections from the dorsal attention network were key in modeling activation differences in the secondary visual network and across all remaining networks. Taken together, these results suggest that shape completion relies upon a distributed but densely interconnected network coalition that is centered in the secondary visual network, coordinated by the dorsal attention network, and inclusive of at least three other networks.

**Highlights:**

- Shape completion differentially activates regions distributed across five networks
- The secondary visual network plays the clearest role in shape completion
- Dorsal attention functional connections likely coordinate activity across networks
- Posterior temporal region, PH, played a highly consistent role in completion

## 1. Introduction

Visual shape completion plays a fundamental role in normal seeing, extracting object shape, size, position, and numerosity from the relative alignments and orientations of spatially segregated edges. Converging evidence from human and non-human primates suggests that the process relies upon V4, LO, V2, and V1, with feedback cascading from the former to the latter two regions. For example, transcranial magnetic stimulation applied earlier to LO (100-122 ms) or later over V1/V2 (160-182 ms) worsened discrimination of completed shapes (Wokke et al., 2013). Multielectrode array recordings of V4 revealed differential activity for completed shapes within ~150 ms, which could plausibly precede low-level visual activations (Cox et al., 2013). In single-cell recordings, deep layer V2 cells responded ~100 ms post-stimulus onset and deep layer V1 cells responded ~120-190 ms (Lee and Nguyen, 2001). In addition to feedback, long-range horizontal excitatory connections between V1 pyramidal cells also bolster edge integration (Iacaruso et al., 2017). These four regions—V1, V2, V4, and LO—have been termed the “classical” regions of shape completion (Keane, 2018) given their inter-connectedness and well-established role in the process.^1^

What other regions participate in shape completion? At present there is no consensus (M. M. Murray and Herrmann, 2013; Seghier and Vuilleumier, 2006). Fusiform gyrus, V3A, and V3B/KO have been implicated (Mendola et al., 1999; M. Murray et al., 2002), although the last region has been found mainly, but not exclusively, with dynamic illusory contour stimuli (Kruggel et al., 2001). In a magnetoencephalography (MEG) study, adults passively viewing briefly-presented pac-man stimuli (30 ms) exhibited more orbitofrontal (OFC) modulation relative to a control stimulus 340 ms post stimulus onset (Halgren et al., 2003). The OFC effect has not been replicated perhaps because older fMRI studies had coarser spatial resolution, more partial voluming, and thus more signal drop-out near the sinuses (due to magnetic field inhomogeneities). Other studies have found activation in the frontal or posterior parietal cortices for illusory “Kanizsa” shapes, but in certain instances there was no control condition or the effects did not eclipse those found for a control condition (Doniger et al., 2002; Foxe et al., 2005; M. M. Murray et al., 2004). A more recent review of illusory contour perception did not mention frontal/prefrontal cortex (M. M.Murray and Herrmann, 2013). Finally, many of the studies that searched for shape completion effects across cortex invoked MEG, EEG, or lower-resolution MRI, and thus had limited ability to locate activations with spatial precision.

Another unanswered question pertains to the functional connections and large-scale networks involved in shape completion. A search of relevant key terms on PubMed retrieved 885 items on shape completion but the list dwindled to zero when either “functional connectivity” or “functional network” was conjoined to the search.^2^ Couching a process in terms of its encompassing network is useful. It allows for a better interpretation of co-modulated regions that fall within that same network. It allows functional interactions to be understood in a larger context and motivates further tests on how the networks interact. Finally, because networks are much larger functional units and much more readily aligned between subjects, network-based results are easier to generalize across subjects (Ji et al., 2019).

There are good reasons to document the neural basis of shape completion. The process is phylogenetically primitive and ontogenetically early, underscoring its importance for normal seeing (Nieder, 2002; Valenza and Bulf, 2010). Moreover, shape completion deficits arise during brain injury (Vuilleumier et al., 2001), developmental agnosia (Gilaie-Dotan et al., 2009), sight restoration (Ostrovsky et al., 2009), and neuropsychiatric illness (Keane et al., 2019). Knowing the neural basis of shape completion constitutes a first step for developing novel pharmacologic or stimulation-based interventions.

We investigated the brain network mechanisms of shape completion with four task scans and one resting-state scan. Our ability to detect effects was augmented by having used a multiband temporal resolution to increase signal-to-noise, a smaller voxel size (2.4 mm) to reduce signal drop-out near the ventral surface, a cortex-wide surface-based analysis to improve anatomical accuracy (Glasser et al., 2013), and a parcellation schema to provide a principled way to segregate cortex into a manageable number of functional units. In the task scans, participants discriminated pac-man configurations that formed or failed to form visually completed shapes (illusory and fragmented condition, respectively) (Ringach and Shapley, 1996). Shape completion was operationalized as the difference in performance or activation between the two conditions. This so-called “fat/thin” task was chosen because it has been extensively investigated via psychophysics, fMRI, EEG, and TMS and because it relies upon the classical brain regions just mentioned (Gold et al., 2000; Keane et al., 2007; Maertens et al., 2008; M. M. Murray et al., 2006; Pillow and Rubin, 2002; Wokke et al., 2013). The resting-state scan data allowed us to compute the resting-state functional connectivity (RSFC) matrix between all pairs of regions, which in turn allowed us to assess the likely utility of the functional connections for shape completion via a recent brain activity mapping procedure dubbed “ActFlow” (Cole et al., 2016). The ActFlow method estimates the actual task activation difference (illusory-fragmented) for a given target region by taking the sum of all other task activation differences (in all other regions) weighted by their functional connectivity strength to that target. If the correlation between actual and estimated activation differences is greater than zero across regions for a subject and if this correlation is significantly above zero across subjects (evaluated via a t-test), then the resting-state connections are likely involved in shape completion. The ActFlow approach is justified since task and rest generate highly similar brain-wide functional connectivity (Cole et al., 2014) and since integrating RSFC into ActFlow has yielded accurate inferences of task-evoked activations in previous studies (Cole et al., 2016).

The results are described in six sections. First, we performed a task activation analysis comparing the task conditions, with careful consideration given to between-task difficulty differences. Second, null V1/V2 effects in the univariate analysis motivated us to perform a post-hoc multivariate pattern analysis (MVPA) to for probe for finer-grained task effects. Third, we divided the parcels into 12 different networks with the Cole-Anticevic Brain Network partition (Ji et al., 2019) and quantified each network’s contribution to shape completion by applying MVPA to parcel-wise task-activations. Fourth, we determined the inter-connectedness of task regions by computing the resting-state functional connectomes (RSFC matrices). Fifth, we demonstrated the likely utility of these functional connections for shape completion via ActFlow; that is, we showed that the task activation difference in each parcel could be inferred from the task activation differences of the remaining parcels weighted by their resting-state connections to that parcel. Finally, again using ActFlow, we determined which network contained the most informative resting-state connections for inferring differential task activity in the secondary visual network (whose relevance was established in Step 3) and across networks. We conclude by suggesting the existence of a shape completion network coalition, which is seated in the secondary visual network, is coordinated by the dorsal attention network, incorporates pieces of three other networks, and interacts with early visual areas at a vertex-wise spatial resolution.

## 2. Materials and Methods

### 2.1. Participants

The sample consisted of healthy controls who participated in a larger clinical study on the neural basis of abnormal visual perceptual organization in schizophrenia and bipolar disorder. These results are thus considered a first step in identifying how the brain represents visually completed shapes in health and disease. (Patient data collection is still ongoing and will be reported once sufficient sample sizes are achieved.) The sample comprised 20 psychophysically naïve participants (2 left handed, 8 females) with an average age of 37.6 and a racial composition of 35% African American, 10% Asian, 35% Caucasian, 15% mixed, and 5% unknown. A quarter of the participants were of Hispanic ethnicity. To obtain a more representative sample, we preferentially recruited controls without four-year college degrees, so that the average number of years of education was 14.8. Written informed consent was obtained from all subjects after explanation of the nature and possible consequences of participation. The study followed the tenets of the Declaration of Helsinki and was approved by the Rutgers Institutional Review Board. All participants received monetary compensation and were naive to the study’s objectives.

The inclusion/exclusion criteria were: (1) age 21-55; (2) no electroconvulsive therapy in the past 8 weeks; (3) no neurological or pervasive developmental disorders; (4) no drug dependence in the last three months (i.e., participants must not have satisfied more than one of the 11 Criterion A symptoms of DSM-5 substance use disorder in the last three months); (5) no positive urine toxicology screen or breathalyzer test on the day of testing; (6) no brain injury due to accident or illness (e.g., stroke or brain tumor); (7) no amblyopia (as assessed by informal observation and self-report); (8) visual acuity of 20/32 or better (with corrective lenses if necessary); (9) the ability to understand English and provide written informed consent; (10) no scanner related contraindications (no claustrophobia, an ability to fit within the scanner bed, and no non-removable ferromagnetic material on or within the body); (11) no DSM-5 diagnosis of past or current psychotic or mood disorders; (12) no current psychotropic- or cognition-enhancing medication; (13) no first-degree relative with schizophrenia, schizoaffective, or bipolar disorder (as indicated by self-report).

### 2.2. Assessments

Psychiatric diagnosis exclusion was assessed with the Structured Clinical Interview for DSM-5 (SCID) (APA, 2000; First et al., 2002). Intellectual functioning of all subjects was assessed with a brief vocabulary test that correlates highly (*r*=0.80) with WAIS-III full-scale IQ scores (Shipley et al., 2009, p. 65; Canivez and Watkins, 2010). Visual acuity was measured with a logarithmic visual acuity chart under fluorescent overhead lighting (viewing distance = 2 meters, lower limit =20/10), and in-house visual acuity correction was used for individuals without appropriate glasses or contacts.

### 2.3. Experimental Design and Statistical Analysis

#### 2.3.1. Stimulus and procedure

Participants performed a “fat/thin” shape discrimination task in which they indicated whether four pac-men formed a fat or thin shape (“illusory” condition) or whether four downward-facing pac-men were uniformly rotated left or right (“fragmented” condition) (see Fig. 1). The fragmented task is a suitable control in that it involves judging the lateral properties of the stimulus—just like the illusory condition—and in that it uses groupable elements (via common orientation, Beck, 1966). Moreover, the two tasks rely on many of the same processes: (1) learning two response alternatives from a limited number of practice exemplars and instructional screens (novel task learning); (2) transferring the learned alternatives to long term memory (consolidation); (3) attending to four discrete spatial regions (divided attention); (4) continuously monitoring the display over specific trial intervals (temporal attention); (5) capturing and extracting spatial information from briefly presented arrays (visual short term memory); (6) discerning fine-grained orientation differences (orientation perception); and (7) repeating the foregoing processes over the task duration (sustained motivation) (Keane et al., 2019). Perhaps because of all these similarities, the two tasks generate similar performance thresholds (Keane et al., 2014) and are highly correlated behaviorally (Keane et al., 2019), which should not be taken for granted being that extremely similar visual tasks are often uncorrelated even with large samples (Grzeczkowski et al., 2017). In sum, by having employed a closely matched and already-tested control condition, we are in a position to identify mechanisms relatively unique to shape completion.

**Fig. 1.**
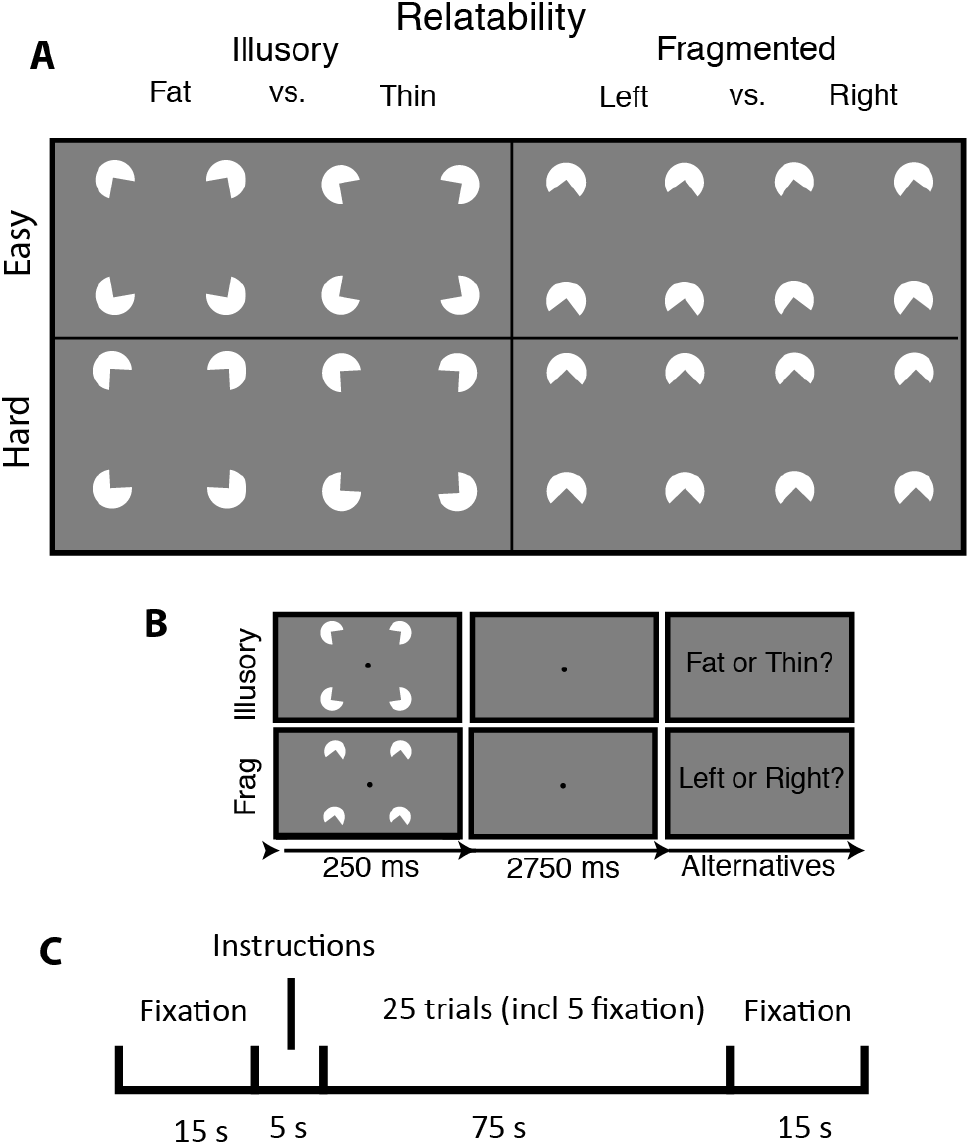
Stimuli, trial sequence, and block arrangement for the visual shape completion experiment. (A) Sectored circles (pac-men) were oriented to generate visually completed shapes (illusory condition) or fragmented configurations that lacked interpolated boundaries (fragmented condition). There were two difficulty conditions corresponding to the amount by which the pac-men were individually rotated to create the response alternatives. (B) After briefly seeing the target, subjects responded. (C) Each half of a run consisted of a fixation screen, a 5 second instructional screen, 25 trials of a single task condition (including 5 fixation trials), and then another fixation screen.

**Fig. 2.**
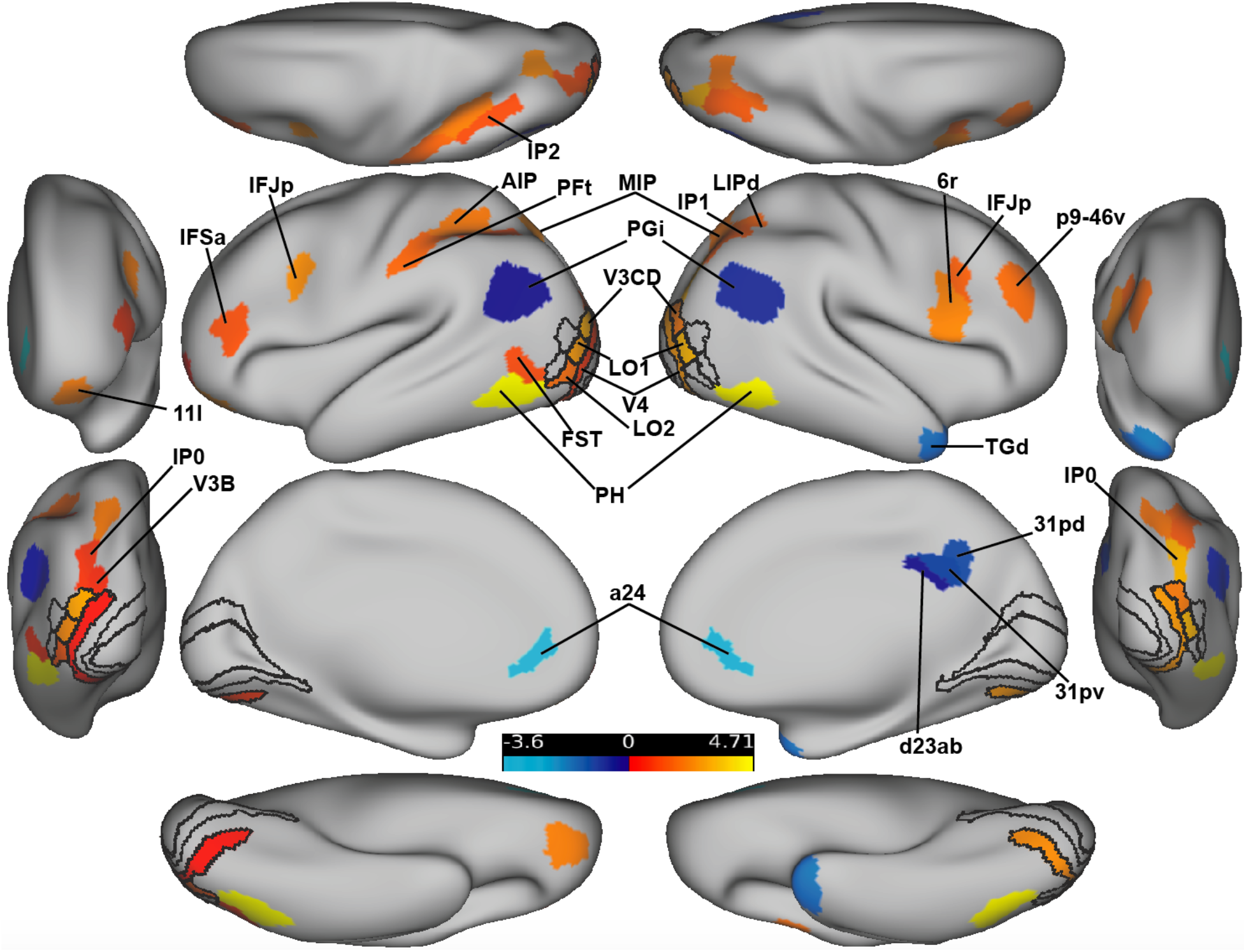
FDR-corrected activation difference amplitudes (Z-normalized) for all parcels for the illusory – fragmented contrast. ROIs are shown with black outlines. The anterior and posterior views are shown laterally; the dorsal and ventral views are shown at the top and bottom. Hot colors indicate regions that were more active for the illusory versus fragmented task; cool colors indicate the reverse.

Subjects viewed the stimuli in the scanner from a distance of 99 cm by way of a mirror attached to the head coil. There were four white sectored circles (radius = .88 deg, or 60 pixels) centered at the vertices of an invisible square (side = 5.3 deg, or 360 pixels), which itself was centered on a gray screen (RGB: 127; see Fig. 3). Stimuli were initially generated with MATLAB and Psychtoolbox code (Pelli, 1997) with anti-aliasing applied for edge artifact removal; images were subsequently presented in the scanner via PsychoPy (version 1.84; (Peirce, 2007) on a MacBook Pro. Illusory contour formation depended on the geometric property of “relatability” (Kellman and Shipley, 1991): when the pac-men were properly aligned (relatable), the illusory contours were present (the “illusory” condition); when misaligned (unrelatable), they were absent (“fragmented” condition).

**Fig 3.**
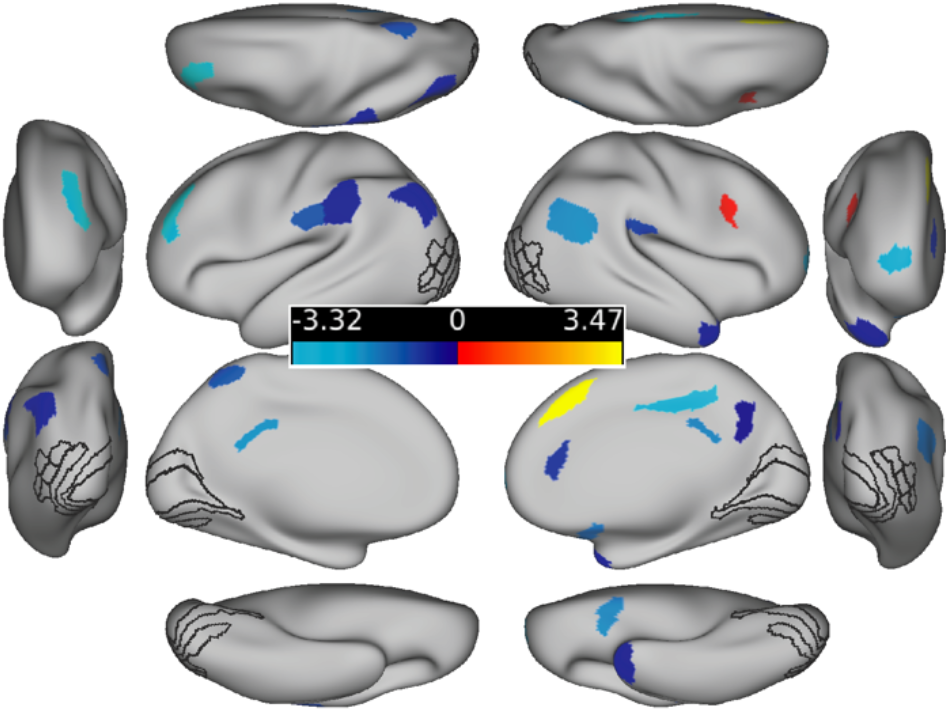
Task activation differences for hard - easy trials (collapsed across illusory/fragmented). Opposite to the illusory-fragmented contrast, we found that harder trials generally elicited less activation throughout the brain relative to easier trials and the location of these significant activations overlapped little with the activations shown in Fig. 2. The illusory/fragmented a priori ROIs (black outlines) are shown for comparison purposes only and did not contain significant parcels.

Within each of the four runs, there was one block of each task condition. In the illusory block, subjects indicated whether four pac-men formed a fat or thin shape; in the fragmented block, subjects indicated whether four downward-facing pac-men were each rotated left or right (see Fig. 1). Block ordering (illusory/fragmented or vice versa) alternated from one run to the next. Each block had two difficulty levels, corresponding to the magnitude of pac-man rotation (+/- 10 degrees “easy”, or +/- 3 degrees of rotation, “hard”). Within each block, there were 20 task trials and 5 fixation trials. Half of the task trials were easy, and half were hard; half of these two trial types were illusory, and half were fragmented. The ordering of these trial types (including fixation) was counterbalanced. Each trial consisted of a 250 ms pac-man stimulus (task trial) or 250 ms fixation dot (fixation trial), followed by a 2750 ms fixation dot. Subjects needed to issue a response before the end of a task trial; otherwise, a randomly selected response was assigned at the end of that trial and the following trial ensued. Feedback was provided at the end of each run in the form of accuracy averaged cumulatively across all test trials.

Subjects received brief practice outside of and within the scanner before the actual experiment. During practice, subjects were reminded orally and in writing to keep focused on a centrally-appearing fixation point for each trial. To ensure that subjects thoroughly understood the task, pictures of the fat/thin stimuli were shown side-by-side and in alternation so that the differences could be clearly envisaged. Subjects issued responses with a two-button response device that was held on their abdomens with their dominant hand; subjects practiced with this same type of device outside of the scanner facility. Feedback after each trial was provided during the practice phase only (“correct”, “incorrect”, or “slow response”).

#### 2.3.2. fMRI acquisition

Data were collected at the Rutgers University Brain Imaging Center (RUBIC) on a Siemens Tim Trio scanner. Whole-brain multiband echo-planar imaging (EPI) acquisitions were collected with a 32-channel head coil with TR = 785 ms, TE = 34.8 ms, flip angle = 55°, bandwidth 1894/Hz/Px, in-plane FoV read = 211 mm, 60 slices, 2.4 mm isotropic voxels, with GRAPPA (PAT=2) and multiband acceleration factor 6. Whole-brain high-resolution T1-weighted and T2-weighted anatomical scans were also collected with 0.8 mm isotropic voxels. Spin echo field maps were collected in both the anterior-to-posterior and posterior-to-anterior directions in accordance with the Human Connectome Project preprocessing pipeline (version 3.25.1) (Glasser et al., 2013). After excluding dummy volumes to allow for steady-state magnetization, each experimental functional scan spanned 3 min and 41 s (281 TRs). Scans were collected consecutively with short breaks in between (subjects did not leave the scanner). An additional 10-minute resting-state scan (765 TRs) occurred in a separate session, with the same pulse sequence. Note that collecting multiband (rather than single-band) data plausibly boosted the signal-to-noise ratio (with its higher spatial resolution) and allowed better detection of structures along the ventral cortical surface (by minimizing partial voluming) (Merboldt et al., 2000; Smith et al., 2013).

#### 2.3.3. fMRI preprocessing

Preprocessing steps are highly similar to earlier studies (Ito et al., 2017) but are repeated below. Imaging data were preprocessed using the publicly available Human Connectome Project minimal preprocessing pipeline which included anatomical reconstruction and segmentation, and EPI reconstruction, segmentation, spatial normalization to standard template, intensity normalization, and motion correction (Glasser et al., 2013). All subsequent preprocessing steps and analyses were conducted on CIFTI 64k grayordinate standard space. This was done for the parcellated time series using the Glasser et al. (2016) atlas (i.e., one BOLD time series for each of the 360 cortical parcels, where each parcel averaged over vertices). The Glasser surface-based cortical parcellation combined multiple neuroimaging modalities (i.e., myelin mapping, cortical thickness, task fMRI, and RSFC) to improve confidence in cortical area assignment. The parcellation thus provides a principled way to parse the cortex into manageable number of functionally meaningful units and thereby reduce the number of statistical comparisons. Note also that there are 97 newly-defined cortical areas in this parcellation, making it possible to identify entirely new shape completion regions. To conduct a follow-up MVPA analysis within V1 and V2 (see Results), we also performed an otherwise identical preprocessing pipeline on the vertex-wise data. In all cases, we performed nuisance regression on the minimally preprocessed task data using 24 motion parameters (6 motion parameter estimates, their derivatives, and the squares of each) and the 4 ventricle and 4 white matter parameters (parameter estimates, the derivates, and the squares of each) (Ciric et al., 2017). For the task scans, global signal regression, motion scrubbing, spatial smoothing, and temporal filtering were not used. Each run was individually demeaned and detrended (2 additional regressors per run).

The resting-state scans were preprocessed in the same way as the parcellated task data (including the absence of global signal regression) except that we removed the first five frames and applied motion scrubbing (Power et al., 2012). That is, whenever the framewise displacement for a particular frame exceeded 0.3 mm, we removed that frame, one prior frame, and two subsequent frames (Schultz et al., 2018). Framewise displacement was calculated as the Euclidean distance of the head position in one frame as compared to the one preceding.

Functional and anatomical scans were visually inspected for quality. In addition, an MRI quality control package (“MRIQC”) and an accompanying random forest classifier were used to confirm that all T1 anatomical scans were artifact free (Esteban et al., 2017). (Two other participants, not included in our analyzed sample, had been flagged by MRIQC as having low quality T1 scans.) The mean framewise displacement across scans before motion correction or scrubbing was remarkably similar in the visual completion and rest scans: 0.142 mm for visual completion (averaged across scans) and 0.143 mm for rest. The average number of frames remaining after scrubbing for the rest scan was 696 [range: 548-760].

For the task scans, there were 6 task regressors, one for each instructional screen (illusory/fragmented) and one for each of the four trial types (illusory/fragmented, easy/hard). A standard fMRI general linear model (GLM) was fit to task-evoked activity convolved with the SPM canonical hemodynamic response function (using the function spm_hrf.m). Betas for the illusory and fragmented condition were derived from all trials of the relevant condition across all four runs. For the classifier analyses, described below, task activation betas were derived separately for each run, but all other steps were the same as described.

#### 2.3.4. Task activation and multivariate pattern analyses

Analyses were performed with RStudio (Version 1.2.1335) and MATLAB R2018b. Cortical visualizations were created with Workbench (version 1.2.3). There were eight parcels of a priori interest in each hemisphere. These ROIs have been given different names in different research studies (shown in parentheses) and are as follows: V1 (17, hOC1, OC, BA17), V2 (18, hOC2, OB, BA18), V4 (V4d, V4v, hV4, hOC4v, hOC4lp), V4t (LO2), LO1 (LO2, hOC4la); LO2 (LO1, hOC4la), LO3 (hOC4la), and V3CD (V3A,V3B, hOC4la) (Glasser et al., 2016, p. 81 see of Supplementary Neuroanatomical Results). Note that V3CD was included because it corresponds to the anterior third of the middle and inferior lateral occipital gyri (area hOc4la as labeled by Malikovic et al., 2016). Statistical correction, when applied, was via the False Discovery Rate (FDR) method (Benjamini and Hochberg, 1995). For the univariate task activation analysis, regions that were and were not of a priori interest were separately FDR-corrected. (Statistical correction is indicated explicitly in the text below, e.g., via p_corr_ values).

For the group-level task activation analyses, betas for each subject were derived for each parcel, averaged across difficulty condition, and subtracted (illusory-fragmented). These values were then compared to zero across subjects with a one-sample t-test. As a control analysis, we did the same as just described, except that we averaged across *task* condition and contrasted the easy/hard conditions. As a further demonstration of the robustness of the univariate results, we performed individual subject parcel-wise task activation analyses for the illusory/fragmented contrast (Table 1), using the subject’s estimated covariance matrix, task betas, and MATLAB’s linear hypothesis test function (linhyptest).

**Table 1.** *Results for parcels that that were either of a priori interest or that were significant on the illusory-fragmented task activation analysis* (see Fig. 2). The rows were sorted in descending order, first, by the percentage of subjects showing the effect in the group direction (column 2) and, then, by the percentage of subjects showing significant effects on the individual subject analysis (column 3). The prefix of each parcel name (“L_ “or “R_”) indicated its hemisphere. The fourth and fifth columns indicate a parcel’s ROI status (yes/no) and functional network. The next three columns indicate whether a parcel was significant after FDR correction, whether it remained significant when task conditions were matched on accuracy/RT, and whether it was significant using the predicted ActFlow data. In the final column, we show the average task activation difference, with more positive values indicating more illusory relative to fragmented activation.

| Parcel Name | % With Difference In Group Direction | % With Sig. Difference | ROI? | Network | Sig. with FDR correction? | Sig. with accuracy matching ? | Sig. with Act-Flow? | Mean Beta Difference [95% CI] |
| --- | --- | --- | --- | --- | --- | --- | --- | --- |
| R_PH | 100 | 70 | 0 | Visual2 | 1 | 1 | 1 | 111.5 [75.6,147.5] |
| L_PH | 95 | 80 | 0 | Visual2 | 1 | 1 | 1 | 93.2 [ 63.7,122.7] |
| L_MIP | 95 | 60 | 0 | Dorsal-attention | 1 | 1 | 1 | 80.8 [ 41.9,119.7] |
| R_V4 | 95 | 55 | 1 | Visual2 | 1 | 1 | 1 | 47.4 [ 25.7, 69.1] |
| L_IFJp | 90 | 50 | 0 | Frontoparietal | 1 | 1 | 1 | 87.5 [ 45.8,129.2] |
| R_p9-46v | 90 | 50 | 0 | Frontoparietal | 1 | 1 | 1 | 60.5 [ 27.4, 93.5] |
| L_LO1 | 90 | 40 | 1 | Visual2 | 1 | 1 | 1 | 66.7 [ 36.4, 97.1] |
| R_a24 | 90 | 35 | 0 | Default | 1 | 1 | 1 | -44.5 [-66.2,-22.9] |
| L_V3CD | 85 | 75 | 1 | Visual2 | 1 | 1 | 1 | 76.1 [ 43.8,108.3] |
| L_PFt | 85 | 60 | 0 | Dorsal-attention | 1 | 1 | 1 | 60.5 [ 26.5, 94.5] |
| R_IP0 | 85 | 60 | 0 | Dorsal-attention | 1 | 1 | 1 | 76.1 [ 44.7,107.6] |
| R_V3CD | 85 | 60 | 1 | Visual2 | 1 | 1 | 1 | 67.8 [ 32.9,102.8] |
| L_V4 | 85 | 55 | 1 | Visual2 | 1 | 0 | 1 | 41.0 [ 13.6, 68.3] |
| L_a24 | 85 | 55 | 0 | Default | 1 | 1 | 1 | -49.5 [-73.1,-25.8] |
| R_LIPd | 85 | 55 | 0 | Dorsal-attention | 1 | 1 | 1 | 78.8 [ 34.6,123.1] |
| R_TGd | 85 | 50 | 0 | Default | 1 | 1 | 1 | -36.4 [-56.9, -16.0] |
| L_IP0 | 85 | 45 | 0 | Dorsal-attention | 1 | 0 | 1 | 64.1 [ 25.3,102.8] |
| R_LO1 | 85 | 45 | 1 | Visual2 | 1 | 1 | 1 | 58.5 [ 34.4, 82.5] |
| R_PGi | 85 | 45 | 0 | Default | 1 | 0 | 1 | -40.0 [-64.5,-15.4] |
| R_IFJp | 85 | 40 | 0 | Frontoparietal | 1 | 0 | 1 | 85.2 [ 36.5,133.8] |
| L_IFSa | 85 | 35 | 0 | Frontoparietal | 1 | 1 | 0 | 48.5 [ 19.3, 77.7] |
| R_31pd | 85 | 35 | 0 | Default | 1 | 0 | 1 | -60.1 [-95.8,-24.4] |
| L_PGi | 85 | 30 | 0 | Default | 1 | 0 | 1 | -36.8 [-60.1,-13.5] |
| L_AIP | 80 | 60 | 0 | Dorsal-attention | 1 | 1 | 1 | 76.7 [ 37.0,116.4] |
| L_11l | 80 | 55 | 0 | Frontoparietal | 1 | 1 | 1 | 60.6 [ 31.8, 89.4] |
| R_MIP | 80 | 55 | 0 | Dorsal-attention | 1 | 1 | 1 | 79.1 [ 37.7,120.5] |
| R_6r | 80 | 55 | 0 | Cingulo-Opercular | 1 | 1 | 1 | 53.0 [ 26.6, 79.5] |
| L_V3B | 80 | 50 | 0 | Visual2 | 1 | 1 | 0 | 52.0 [ 19.5, 84.5] |
| L_LO2 | 80 | 45 | 1 | Visual2 | 1 | 1 | 1 | 50.4 [ 23.1, 77.7] |
| R_LO2 | 80 | 45 | 1 | Visual2 | 0 | 0 | 1 | 47.3 [ 5.7, 88.8] |
| L_FST | 80 | 35 | 0 | Visual2 | 1 | 0 | 1 | 46.7 [ 17.8, 75.5] |
| L_IP2 | 75 | 60 | 0 | Frontoparietal | 1 | 1 | 1 | 70.9 [ 27.0,114.7] |
| R_d23ab | 75 | 30 | 0 | Default | 1 | 0 | 1 | -53.9 [-88.1,-19.7] |
| R_IP1 | 70 | 55 | 0 | Frontoparietal | 1 | 0 | 0 | 52.1 [ 21.8, 82.4] |
| L_LO3 | 70 | 35 | 1 | Visual2 | 0 | 0 | 1 | 26.6 [-12.3, 65.6] |
| R_31pv | 70 | 30 | 0 | Default | 1 | 0 | 1 | -54.8 [-87.7,-21.9] |
| R_LO3 | 70 | 30 | 1 | Visual2 | 0 | 0 | 0 | 22.9 [-13.5, 59.4] |
| R_V4t | 70 | 25 | 1 | Visual2 | 0 | 0 | 0 | 16.8 [-21.2, 54.9] |
| R_V2 | 60 | 25 | 1 | Visual2 | 0 | 0 | 0 | 3.2 [-30.0, 36.4] |
| L_V2 | 60 | 20 | 1 | Visual2 | 0 | 0 | 0 | 7.0 [-23.4, 37.3] |
| L_V1 | 55 | 30 | 1 | Visual1 | 0 | 0 | 0 | 9.1 [-25.4, 43.7] |
| R_V1 | 55 | 25 | 1 | Visual1 | 0 | 0 | 0 | 7.0 [-28.3, 42.2] |
| L_V4t | 50 | 15 | 1 | Visual2 | 0 | 0 | 0 | -0.2 [-34.9, 34.5] |

The location and role of each parcel was considered within the context of their functional network affiliations. We used the Cole-Anticevic Brain Network partition, which comprised 12 functional networks that were constructed from the above-mentioned parcels and that were defined via a General Louvain community detection algorithm using resting-state data from 337 healthy adults (Ji et al., 2019 see Fig. 4A). This partition included well-known sensory networks—primary visual, secondary visual, auditory, somatosensory; previously identified cognitive networks—frontoparietal, dorsal attention, cingulo-opercular, and default mode; a left-lateralized language network; and three entirely novel networks—posterior multimodal, ventral multimodal, and orbito-affective. This partition passed several quality control measures of stability and reliability, was biologically motivated and statistically principled, and was able to demonstrate increased levels of network-level task activations.

**Fig. 4.**
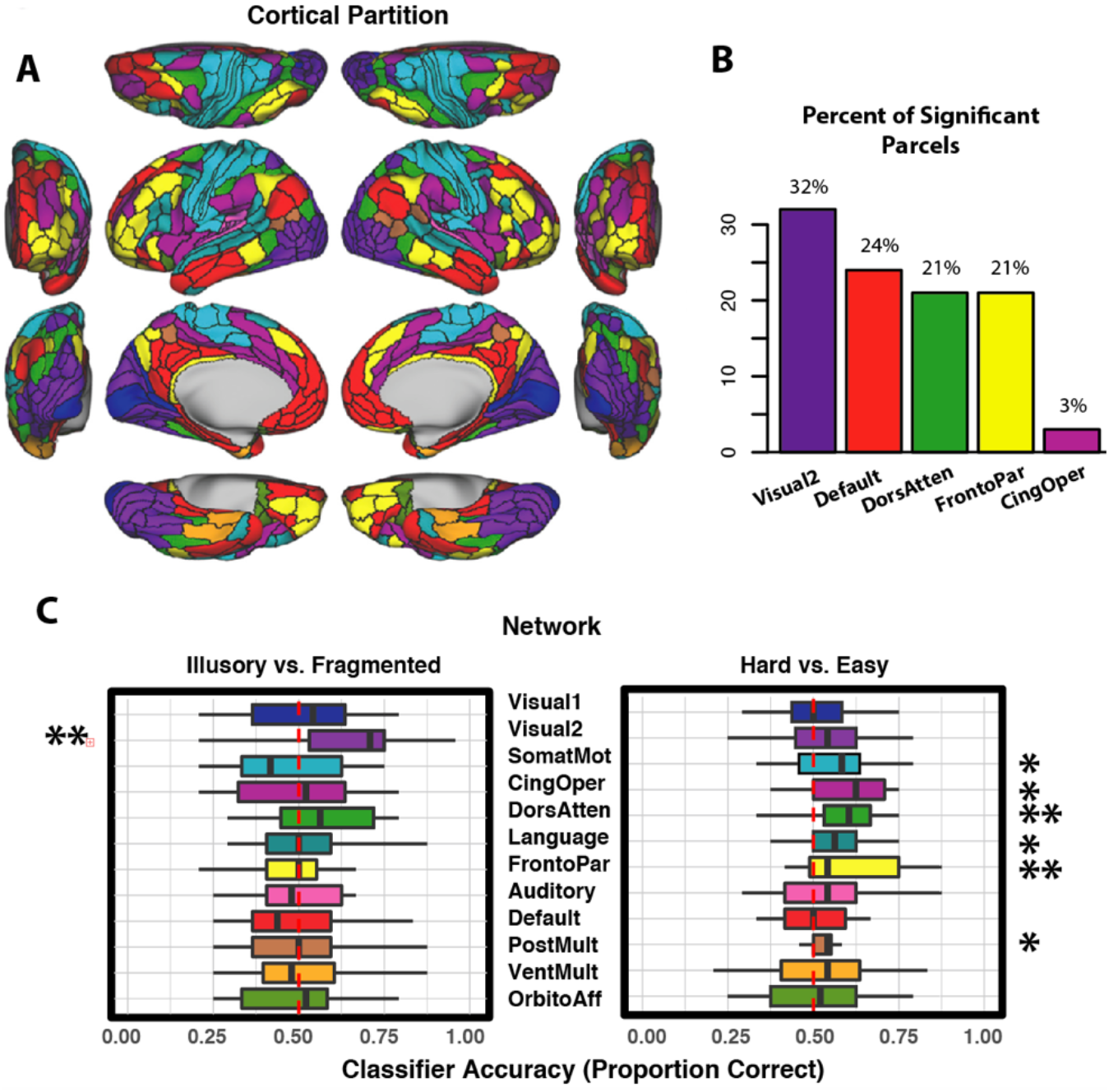
(A) The Cole-Anticevic Brain Network partition. We considered whether parcel-wise activation patterns in the cortical networks could individually classify task betas as deriving from the illusory or fragmented condition; these included the primary visual, secondary visual, somatomotor, cingulo-opercular, dorsal attention, language, frontoparietal, auditory, default, posterior multimodal, ventral multimodal, and orbito-affective networks. Networks are color coded to match the parcels in panels B and C. (B) The percentage of significantly modulated parcels that belonged to each network for the illusory/fragmented contrast. (C) Classification accuracy for the illusory/fragmented and hard/easy comparisons. The red dotted line shows chance performance, the box segments denote median scores, the box hinges correspond to the 25th and 75th percentiles, and the box whiskers extend to the largest or smallest value (but no further than 1.5x the interquartile range). Only the secondary visual network could significantly predict illusory/fragmented activations. In comparison, multiple networks were involved in classifying easier/harder trials, after FDR correction (*p_corr_<.05, ** p_corr_<.01).

Multivariate pattern analysis was performed on the activation betas at two levels of spatial granularity. First, we examined whether 12 different functional networks could individually classify task condition (illusory vs fragmented) or difficulty condition (easy vs hard) using their within-network mean parcel activations as features. Next, on a follow-up post-hoc analysis, we examined, for each parcel, whether vertex-wise activations could classify task condition. MVPA classification accuracy in each case was assessed via leave-two-runs-out cross validation. For example, when classifying task condition for each participant, we examined whether the betas for each of the two left-out runs better correlated to the averaged illusory or fragmented betas of the remaining runs. Note that each run contained an equal number of trials from each of the two conditions, ensuring balanced condition types across test and training. Pearson correlation served as the minimum distance classifier (i.e., 1-r) (Mur et al., 2009; Spronk et al., 2018). Results were averaged for each subject across the 6 possible ways to divide the four runs between test and validation. Statistical significance was determined via permutation testing, which generated a null distribution of classification accuracies through the same procedure with 10,000 samples. That is, for each sample, the “illusory” and “fragmented” labels were shuffled for each subject and run, and the classification results were averaged across subjects and across the 6 possible divisions of testing and validation data sets.

#### 2.3.5. Resting-state functional connectivity derivation

We determined the resting-state functional connections for each parcel. Specifically, for each target parcel time series, we decomposed the time series of the remaining (N=359) parcels into 100 components, regressed the target onto the PCA scores, and back-transformed the PCA betas into a parcel-wise vector. The average amount of variance explained by the components across subjects was 84% [range: 81-88%]. The RSFC computation is equivalent to running a multiple regression for each parcel, with all other parcels serving as regressors. An advantage of using multiple regression is that it removes indirect connections (Cole et al., 2016). For example, if there exists a true connection from A to B and B to C, a Pearson correlation, but not regression, would incorrectly show connections between A and C. PC regression was preferred over ordinary least squares to prevent over-fitting (using all components would inevitably capture noise in the data). The RSFC matrix served two functions. First, it obviously allowed us to ascertain the functional connectedness of modulated task regions. Second, it allowed an assessment of the utility of these connections for estimating task activation differences via ActFlow.

#### 2.3.6. Activity flow mapping and measuring out-of-network contributions

Fig. 6 illustrates how we used resting-state data to predict task activation (“Activity Flow mapping” or simply “ActFlow”), where the “activations” in this case correspond to the illusory-fragmented difference. For each subject, the task activation in a held-out parcel (‘j’ in Fig. 6A) was predicted as the weighted average of the activations of all other parcels, with the weights being given by the resting-state connections. That is, for each subject, each held out region’s predicted value was given as the dot product of the task activations in the remaining regions (‘i’ in Fig. 6A) and the subject’s restFC between j and i (using the FC weight from the appropriately oriented regression, i.e., j as the target and i as the predictor). The accuracy of the activity flow predictions was then assessed by computing the overlap (Pearson correlation) between the predicted and actual task activation difference vector. Overlap can be expressed by comparing actual and predicted activations for each subject, and then averaging the resulting Fisher-transformed r values (r_z_) across subjects (subject-level overlap). Statistical significance can be determined by comparing the vector of r_z_ values to zero via a one-sample t-test. Overlap can also be expressed by averaging the predicted values across subjects and then comparing that to the averaged actual values, which will yield a single Pearson r value (group-level overlap). If the RSFC matrix can be used to predict task activation differences, that would show that those same functional connections likely subserve task performance. Below, we applied ActFlow once to the full RSFC matrix and once to the matrix involving the task modulated regions.

**Fig. 5.**
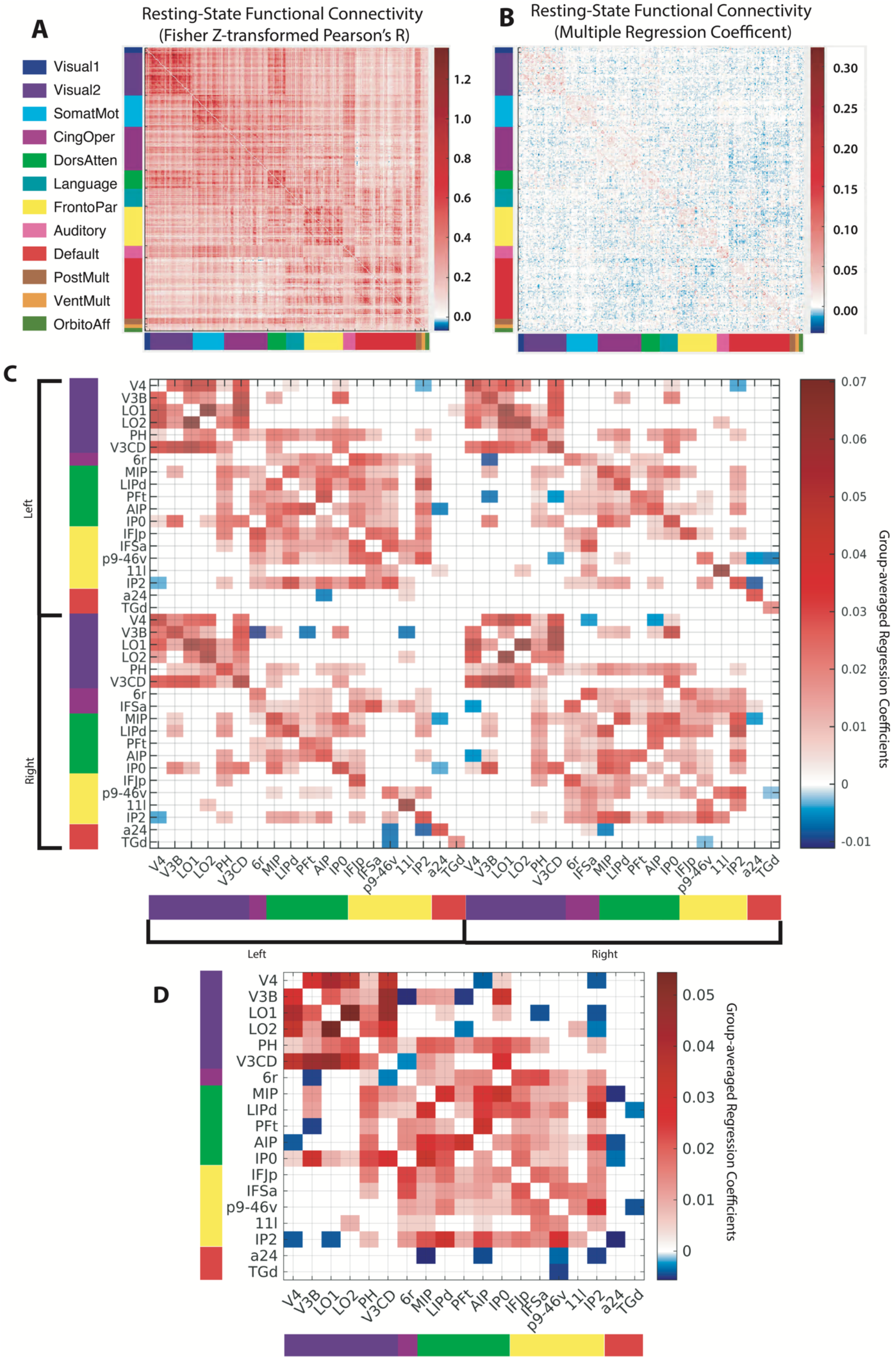
Resting-state functional connectivity (RSFC) matrices. (A) Pearson correlation between the resting-state time series of all parcel pairs (360 x 360 parcels). Parcels are sorted into previously established (color-coded) functional connectivity networks (Ji et al., 2019) (see also Fig. 4A). The block-like structure along the diagonal exemplifies the stronger connectivity within relative to between each network. (B) An RSFC matrix computed via multiple regression (see Methods). The blue/red colors indicate the degree to which a given parcel time series was predicted by all remaining parcels. Note that this matrix is much sparser than the correlational matrix since it eliminates many of the indirect connections between parcels (Cole et al., 2016). (C) Thresholded (FDR-corrected) resting-state connections between significantly modulated task regions (see text), which are ordered first by hemisphere and then by network. Compared to the full matrix in panel B, this pared down matrix had about 1 percent the number of possible connections (matrix elements) and triple the proportion of (FDR-corrected) significant connections. (D) Averaging the connection weights across hemisphere increased the proportion even further (from 40% to 53%), highlighting the broadly symmetric connectivity patterns. Note that one parcel, IFSa, was split between the frontoparietal (left hemisphere) and cingulo-opercular networks (right), and was assigned to the frontoparietal network in this plot since only the frontoparietal parcel was significant in the task activation analysis.

**Fig. 6.**
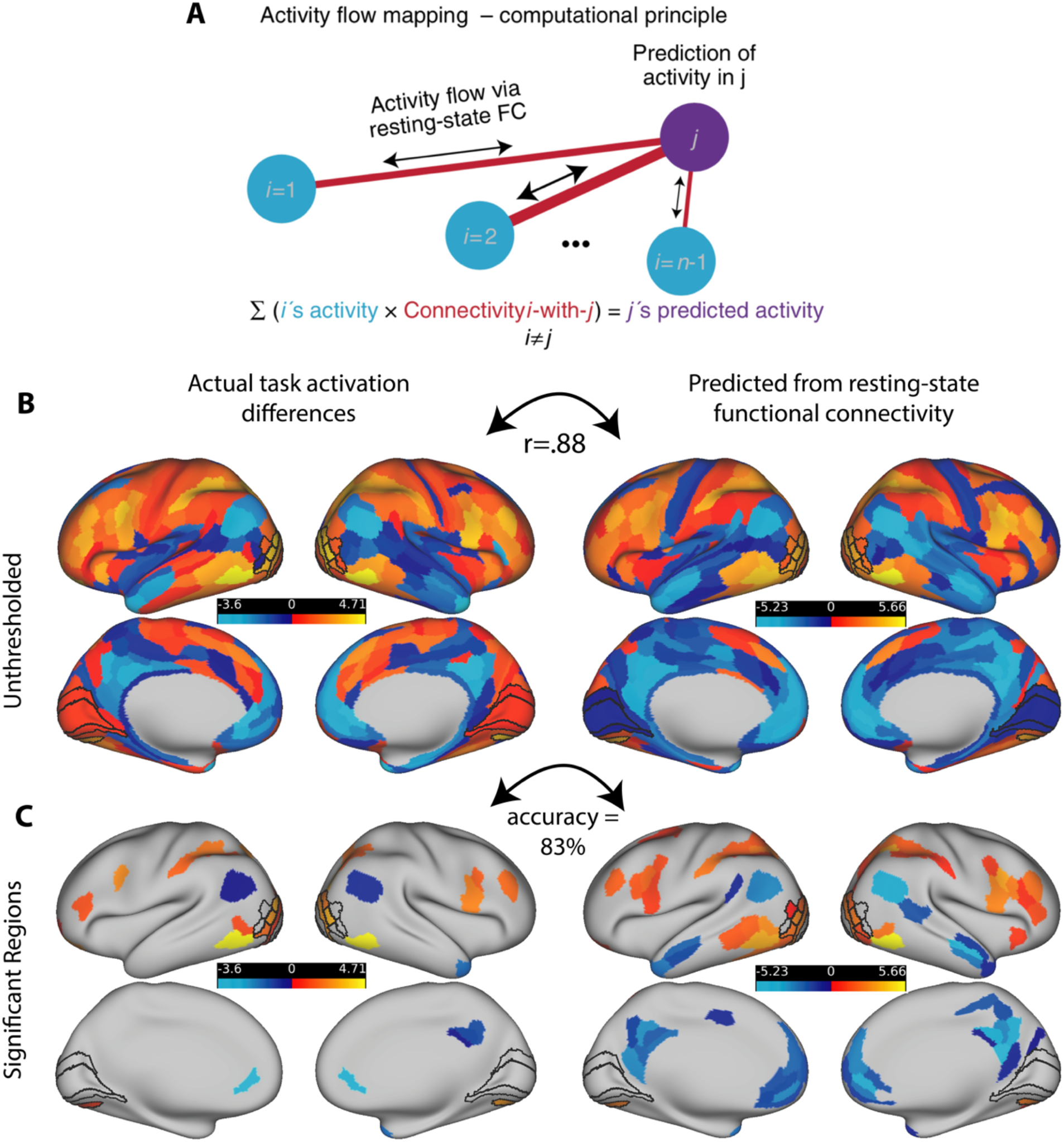
Activity flow mapping for visual shape completion. (A) For each subject, the task activation differences (illusory-fragmented) in a held-out parcel (j) is given by the dot product between the activation differences in the remaining parcels (regions i) and the resting-state connection strengths (betas) between i and j. (B) Unthresholded z-normalized activation differences (illusory – fragmented) as compared to those that were predicted via ActFlow using resting state. (C) When a task activation analysis was applied to the data predicted from ActFlow, statistical significance (or lack thereof) was correctly determined for 83% of the 360 parcels (see also Fig. 2). This suggests that the connection weights derived from resting state are reflective of the actual connections used during shape completion.

Since the secondary visual network was central to the shape completion network coalition, we also examined how ActFlow estimates improved in that network specifically, when any of the remaining four networks were individually added (Fig. 7). This change was ascertained simply by comparing via a paired t-test the prediction accuracies (correlations) before and after adding the network. A significant improvement would indicate which other networks, if any, are important for guiding activity flow in the secondary visual network. The success of the ActFlow method also prompted us to also consider whether adding connections from any of the five task modulated networks could improve ActFlow accuracy in the remaining networks. A significant improvement would indicate which other network, if any, explains differential activity in the remaining networks.

**Fig. 7.**
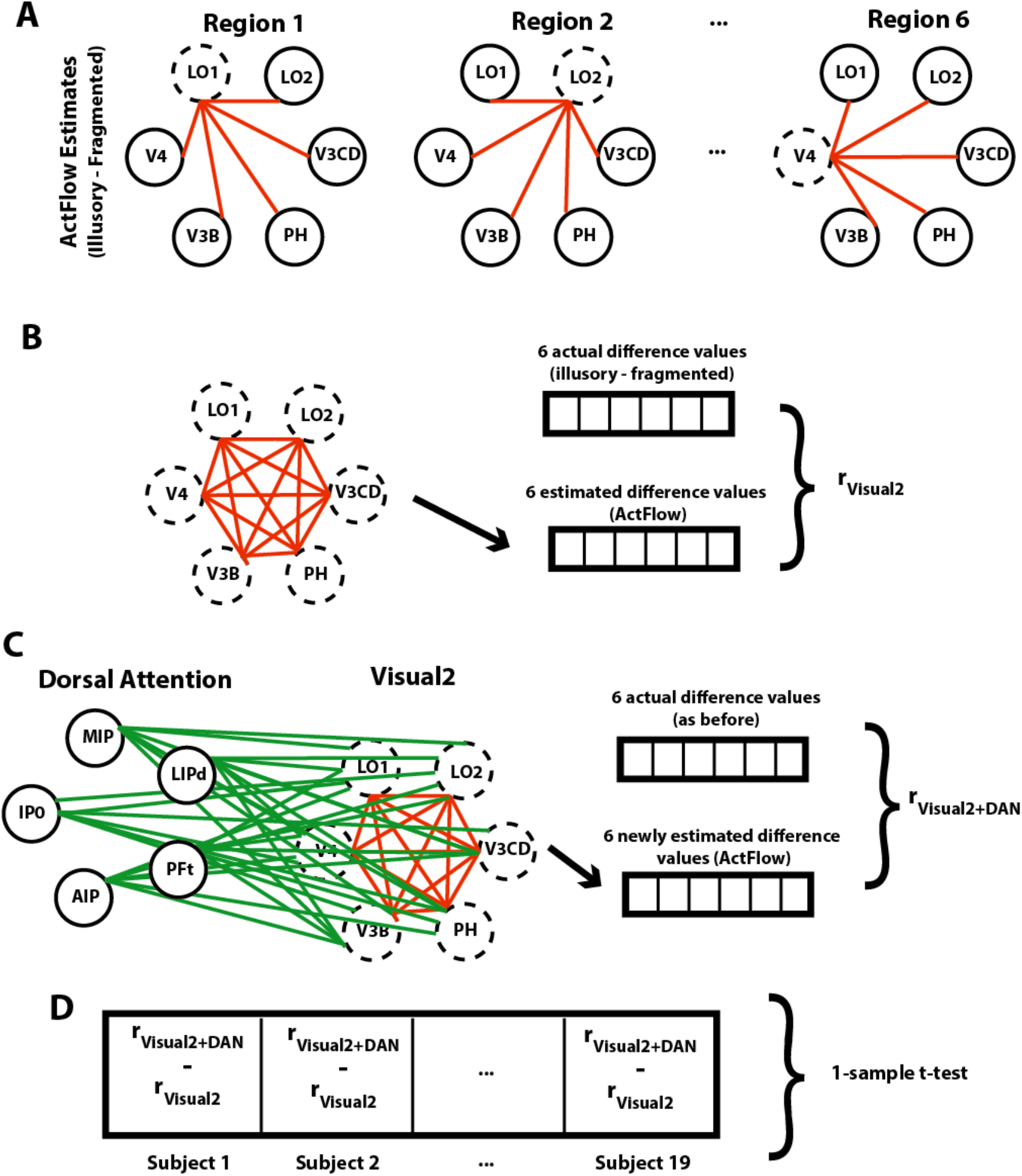
Gauging contributions of the dorsal attention network to the secondary visual network (Visual2). (A) For a given subject, the task activation difference for each significant Visual2 parcel was estimated (dotted circles) using *actual* task activation differences in the remaining parcels (solid circles) and their resting-state connections (red lines). For illustration purposes, only one hemisphere is shown. (B) ActFlow accuracy was defined as the correlation between actual and estimated task activation differences, across the Visual2 parcels. (C) Task activation differences were again estimated via ActFlow, except that, this time, the connections and activation differences from the significant dorsal attention regions could also contribute. (D) The difference between the original and re-calculated estimates was computed for each subject (after a Fisher Z-transform) and compared to zero across subjects. Only the dorsal attention network could significantly improve ActFlow estimates in the secondary visual network.

## 3. Results

### 3.1. Behavioral task performance

Employing a 2 (task condition) by 2 (difficulty) within-subjects ANOVA (type III sum of squares), we found that performance was better in the fragmented than illusory condition (89.6% versus 82.9%, *F*(1,19)=14.8, *p*<.01) and better in the (“easy”) large-rotation condition compared to the “hard” small-rotation condition (*F*(1,19)=133, *p*<10^-9^) (Supplementary Fig. S1). The accuracy difference between illusory and fragmented conditions did not depend on difficulty level, although there was a trend toward a greater difference on the smaller rotation condition (two-way interaction: (*F*(1,19)=3.6, *p*=.07). The marginal interaction probably arose from ceiling effects for the fragmented condition since there was no corresponding interaction in the reaction time data (*F*(1,19)=.14, *p*=.7). Reaction time data where in other ways entirely predictable from the accuracy results, with faster performance in the fragmented than the illusory condition (*F*(1,19)=5.1, *p*=.04), and faster performance in the easy than the hard condition (*F*(1,19)=21.3, *p*<.001). The no-response trials were infrequent, occurring on only 5.5% of the trials on average. The frequency of no-response trials did not vary with difficulty level or task condition nor was there an interaction between difficulty and task condition (*ps*>.25). Consistent with past results (Keane et al., 2019), the fragmented and illusory conditions were highly correlated (accuracy—*r*=.74, *p*<.001; RT—*r*=.81, *p*<.0001), confirming that they were reliant upon a common core of mechanisms. (See Supplementary Fig. S1 for graphical depiction of behavioral results.) The correlations were robust and remained significant when calculated with non-parametric tests or after log-transforming the RT data.

### 3.2. Shape completion effects across five large-scale functional networks

A general linear model task activation analysis determined the parcels that were differentially active in the illusory versus fragmented condition. Overall, 34 parcels reached significance in five different networks (Table 1; Fig. 2). Of these parcels, 26 (76 percent) were more activated for illusory relative to fragmented trials. A priori ROIs, when significant, were all more active relative to the control condition; these include bilateral V3CD, V4, L01, and left L02. These effects were robust and would also be significant if we simply performed a cortex-wide FDR correction. Notable null results were V2 and V1 which will be discussed further below. Additional positively and significantly activated regions resided in the posterior parietal, dorsolateral prefrontal, and orbitofrontal regions; they belonged primarily to the secondary visual, dorsal attention, and frontoparietal networks. All 8 of the regions that were negatively activated in the illusory-fragmented contrast belonged to the default mode network. Note that this finding reflects this network’s established on-task deactivation profile (Anticevic et al., 2012), i.e. greater deactivation for the illusory relative to the fragmented condition, consistent with greater task engagement in the illusory condition.

Because task difficulty was greater in the illusory task, perhaps task difficulty, rather than shape completion, drove the effects just described. We addressed this concern in two ways. First, we performed a contrast comparing activation in the easy versus hard trials, averaged across task conditions. To make the results comparable to before, FDR correction was applied separately to regions that were and were not ROIs. We found 17 parcels that were differentially active, but only four overlapped with the illusory-fragmented contrast (see Fig. 3). Three of these parcels were less active in both the hard-easy and illusory-fragmented comparisons: right d23ab, right TGd, and right PGi; one was more active in both (right IFJp). None of the 16 ROIs of visual shape completion were related to task difficulty. Thus, on this analysis, while the above-mentioned four parcels were confounded with task difficulty, the remaining 30 significant parcels in the illusory/fragmented comparison were not confounded.

To further assess the extent to which task difficulty might account for the aforementioned shape completion effects, we ran an additional analysis that was restricted to the 10 participants who did the best in the illusory relative to the fragmented condition, so that there was no longer an accuracy difference (*t*(9)=-0.443, *p*=0.669, Mean difference in proportion correct=-0.011). In this sample, there was also no reaction time difference between task conditions (*t*(9)=1.63, mean RT difference=-.06 seconds, *p*=.14)). As shown in Table 1, with the exception of left V4, the ROIs that were significant in the earlier analysis remained significant in this restricted sample; these include L01 and V3CD in each hemisphere, left LO2, and right V4 (all *p*<.05, uncorrected). Of the four regions that were significant on both the hard/easy and illusory/fragmented contrast (right d23ab, right PGi, right TGd, right IFJp), only right TGd remained significant and thus is more plausibly independent of task difficulty. The foregoing results were about the same if either 9 or 11 participants were included in this accuracy-matched analysis (See Supplementary materials). Other regions that continued to be significant on this accuracy-matched analysis are shown in Table 1.

To examine the robustness of the task activation effects, we additionally report the percentage of subjects showing significant effects (illusory-fragmented) in the group direction on an individual subject analysis (with a linear hypothesis test, see Methods). This was done for regions that were significant on the task activation analysis as well as for other regions that were of a priori interest. As can be seen from Table 1, about 80% of parcels (ranging from 70-100%, depending on the parcel) showed activation differences in the group direction and about half (35 - 80%) showed effects that were statistically significant. Intriguingly, the posterior temporal region PH–which was not of *a priori* interest—was *most* associated with shape completion, with 80% of subjects showing a significant effect in the left hemisphere and 70% in the right hemisphere, and with at least 95% showing group differences in each hemisphere in the group direction. This region’s surprising role in shape completion is discussed further below.

### 3.3. Fine-grained multivariate traces of shape completion in early visual cortex

Task activation analyses did not reveal shape completion effects in V1 or V2. Because a region could conceivably encode a completed shape in its vertex-wise pattern rather than in its univariate mean (Haynes, 2015), we performed MVPA on vertices within these parcels. For completeness, we considered effects within all 360 parcels. The following were significant: L_V4 (*p*=.02, accuracy=58%), L_LO2 (*p*=.03, accuracy =56%), L_V3CD (*p*=.02, accuracy=58%), R_V1 (*p*=.03, accuracy=56%), R_V4 (*p*=.01, accuracy=57%), and R_LO2 (*p*=.03, accuracy=56%). These effects were not corrected for multiple comparisons but are credible given the strong prior evidence for their involvement (see Introduction). Outside of the ROIs, the only region that was significant after FDR correction was R_PGp (*p*=.03, accuracy=64%). Given the hemispherically similar task activations and the bilateral stimulus displays, we performed the same analysis as above, except that vertices were aggregated (without averaging) across hemisphere to increase sensitivity. The effects were similar to before with effects for: V1 (*p*=.027, accuracy=57%), V4 (*p*=.014, accuracy=58%), LO2 (*p*=.01, accuracy=58%), and LO3 (*p*=.03, accuracy=56%). For regions that were not of a priori interest, the following reached significance after FDR correction: PGp (p_corr_<.0001, accuracy=62.9%) in the secondary visual network and IP1 in the frontoparietal network (p_corr_=.04, accuracy=61%). In sum, V1 but not V2 exhibited modest vertex-wise shape completion effects; additional ROIs (V4, LO2) and a new region, PGp, were also consistently significant on this analysis. Possible reasons for null effects in V2 are considered in the Discussion.

### 3.4. A dominant role for the secondary visual network in shape completion

As shown in Fig. 4, most significant parcels resided in the secondary visual network, followed by the default mode, dorsal attention, frontoparietal, and cingulo-opercular networks. To better quantify the network contributions and compare them to one another, we trained MVPA classifiers separately for the 12 functional networks (Ji et al., 2019), using parcel-wise activations as features (see Methods). After FDR correction (across tests for the 12 networks), the secondary visual network could reliably distinguish the illusory and fragmented conditions (p_corr_=.004, accuracy=63%), but no other network could do so (all p_corr_>.24). Paired t-tests showed that, after FDR correction, the secondary visual network was marginally more predictive than 8 of the remaining 11 networks (all p_corr_<.10). Note that there was no correlation between network classification accuracy and parcel count (*r*=.001, *p*=.997), suggesting that smaller networks were not unduly handicapped.

To assess whether the classification success of the secondary visual network was specific to shape completion and also whether the other networks could be predictive under different circumstances, we additionally ran network-level classifications that distinguished between easy and hard trials. Six networks came out as significant: somatomotor (p_corr_=.03, accuracy=56%), cingulo-opercular (p_corr_=.005, accuracy=60%), dorsal attention (p_corr_=.007, accuracy=57%), language (p_cor_r=.03, accuracy=56%), frontoparietal (p_corr_=.004, accuracy=60%), and posterior multimodal (p_corr_=.03, accuracy=56%). Neither of the visual networks were significant. Thus, our data set and analytic approach could reveal significant effects for a number of networks, but only the secondary visual was relevant when examining visual shape completion. Taken together, these results suggest that—from the perspective of parcel-wise task activation differences—the secondary visual network played a robust, specific, and outsized role in shape completion.

### 3.5. Modulated task parcels were densely inter-connected during rest and were straddled by the dorsal attention network

To determine how modulated task regions were functionally interconnected, we derived a whole-cortex RSFC with Pearson correlation (which is more commonly reported) and then with multiple regression (Fig. 5A,B; see Methods). We then homed in on the significant task regions that remained significant when the illusory/fragmented conditions were matched on accuracy/RT. Since the task activations were hemispherically symmetric, contralateral homologues were included so that there was a 38 x 38 RSFC matrix. The predicted betas from the regression-based RSFC matrix were compared to zero for each connection across subjects (one sample t-test) and were FDR-corrected (thresholded) across all connections. Informal observation of Fig. 5C shows that parcels had higher within-than between-hemisphere RSFC, more cross-hemisphere connections for sensory (visual) than for non-sensory networks, and higher RSFC with their contralateral homologues than with other contralateral regions. These results are consistent with past work (Power et al., 2011; Stark et al., 2008) and demonstrate that the RSFC matrices were yielding sensible results.

A major question was whether the significantly modulated task regions were inter-connected during rest. After applying FDR corrections to each matrix separately, we found that the restricted RSFC matrix (38 x 38) contained three times as many significant resting-state connections as the full (360 x 360) matrix, (39.6 % versus 13.5%). To put this in perspective, two of the twelve resting-state networks—default mode and orbitoaffective—had a lower proportion of significant *within*-network connections (35% and 25%, respectively). This suggests that the significant task regions, despite being composed of five different networks, composed a densely inter-connected network coalition or supra-network. Note that these five networks, as a whole, were not unusually connected to one another: If all regions from all 5 networks were included in the above calculations (to form a 260 x 260 RSFC matrix), the total number of significant resting state connections would still only be 17%. Thus, it is the specific regions within these five networks that appear to be more interconnected during rest.

To examine these results in a different way, we examined for each subject the average within-network connection weight and the average out-of-network connection weight across the 38 task parcels (where “network” consisted of just these parcels), and simply compared these two averaged weights across subjects. Shape completion regions cohered more strongly with one another than with other regions (*t*(19)=19.3, *p*<10^-12^, *d*=3.8).

The RSFC matrices offers clues as to how the regions were communicating. As can be observed from Fig. 5D, the secondary visual network most often connected to the dorsal attention network regions, which in turn had the most significant out-of-network connections (117 connections). Moreover, there appear to be a number of routes between frontal cortex and the mid-level vision ROIs. Dorsal lateral prefrontal cortex (p9-46v) connects with MIP, IPO, and IP2 (in posterior parietal cortex), which in turn connect with all of the significant ROI regions. Intriguingly, area 11l (OFC) connected directly with area LO2. Hence there exist clear routes for conceptual or value-laden information to loop back into areas most typically associated with visual shape completion, but in most cases these routes must traverse the dorsal attention network and particularly parts of posterior parietal cortex.

### 3.6. Resting-state connections are relevant for visual shape completion

We have shown that regions that were differentially activated during visual shape completion were also connected during rest. However, despite some indirect evidence from other work (see Introduction), it remains unclear whether these connections in these same subjects played a mechanistic role in shape completion. To address the question, we leveraged a recently-developed predictive modeling approach—activity flow mapping (“ActFlow”)—to assess whether the resting-state connections (derived via multiple regression) were likely instrumental in carrying the flow of activity between regions during task performance (Cole et al., 2016). In this method, the activation difference (illusory minus fragmented) in a held-out “target” parcel was computed as the linear weighted sum of the activation differences in all other parcels, with the weights being given by the resting-state connections to the target (see Fig. 6A). This can be thought of as a rough simulation of the movement of task-evoked activity between brain regions that likely contributed to each brain region’s task-evoked activity level. This allowed us to assess whether the observed resting-state connections mechanistically supported the perceptual processes associated with shape completion. Prediction accuracy was gauged as the correlation between the actual and predicted activation differences. As can be seen in Fig. 6B, the predictions were highly significant at the whole-cortex level (*r*=.62, *p*<10^-9^). If we were to first average the predicted differences across subjects, then average the actual differences across subjects, and then correlate the two, the resulting group-level accuracy estimate would increase (*r*=.88), probably by increasing the signal-to-noise ratio (Cole et al., 2016).

We next applied a task activation analysis to the ActFlow predicted data (via one-sample t-tests, as before) and compared the results to the original task activation results (shown in Fig. 2). The percentage of parcels that remained significant (sensitivity) with ActFlow was 92%; the percentage of non-significant parcels that remained non-significant (specificity) was 82% (see Fig. 6C). These results again suggest that the observed resting-state connections describe the routes over which task-evoked activity flows during shape completion (controlling for orientation judgement).

To assess the relevance of resting-state connections between regions that were modulated during the task, we restricted activity flow mapping only to those regions and their contralateral homologues. To minimize the chance of task difficulty effects, we again used only regions that remained significant when conditions were matched on accuracy/RT so that each held-out parcel’s activation was predicted by 37 other connections/parcels. Despite eliminating 90 percent of the connections for each parcel, the prediction accuracy estimates (r-values) across subjects were still high (illusory-fragmented: *r*=.58, *p*=5.8*10^-8^) and did not significantly differ (*p*=.36) from the ActFlow correlations with the full matrix (as assessed with a paired t-test). This suggests that much of shape completion can be understood solely by examining the connections and activations of task modulated regions.

### 3.7. Dorsal attention regions can model activity flow in the secondary visual network and across all other networks

According to the task activation and network-wise MVPA results (Table 1 and Fig. 4, respectively), shape completion was most undergirded by the secondary visual network. To examine which other networks might plausibly contribute to the illusory/fragmented activation differences in this network, we determined which ones could improve the ActFlow predictions, using the same significant task regions as before (see Fig. 6). More explicitly, for each subject, we computed a single correlation between the actual and ActFlow parcel difference values across the 12 significant secondary visual network parcels. We then recomputed this correlation, when each of the 12 parcels could *also* be predicted by parcels and connections from one other network. Finally, we Fisher-z transformed the correlations, subtracted the two, and then performed a one-way t-test to see if the correlations increased as a result of the network’s inclusion. The dorsal attention network improved the predictions for the secondary visual network (Δ*r*≈ΔrZ =.33, p_corr_=.01); no other network generated an effect.

Is there a particular network that plays a dominant role in orchestrating the activity of the other regions? We examined this possibility by using the same approach as just described; that is we calculated, for each subject, the ActFlow accuracy for all regions outside of a held-out network and considered how that accuracy improved—that is, how the Fisher-Z correlations increased (ΔrZ)—when the held-out network regions were allowed to contribute (Mill et al., 2020). This was done for each of the five networks, using only the significant task regions (viz., 38 regions were treated as targets for ActFlow in Fig 6A). Consistent with observations from the functional connectivity matrix, the dorsal attention network’s contributions significantly improved predictions for the significant regions of all four remaining networks (Δ*r*≈ΔrZ=.13; *t*(18)=4.84, p_corr_=.0007). Interestingly, every other network—including the secondary visual—failed to influence the results on this analysis (all p_corr_>.17). The improvement from the dorsal attention regions was significant also if we were to use all 360 regions and all possible resting-state connections (rather than restricting to the significantly activated regions; Δ*r*≈ΔrZ=.03; *t*(18)=3.60, p_corr_=.007).

## 4. Discussion

Visual shape completion plays a critical role in extracting object shape, size, position, and number from edge elements dispersed across the field of view. The process relies on lateral occipital and early visual areas, but it is unclear what other regions might be utilized, how they are functionally connected, or what networks they reside within. To shed light on the foregoing, we scanned participants during rest and during a task in which they discriminated pac-man quartets that either formed or failed to form visually completed shapes. Six major findings emerged. One is that although only a few dozen parcels were differentially activated, the effects were impressively consistent, with one region—parcel PH—exhibiting similar effects across 95% of subjects in each hemisphere. Next, the secondary visual network played a dominant role in shape completion but parcels within the dorsal attention, frontoparietal, and default mode, and cingulo-opercular network were also influential, suggesting that shape completion is a distributed process. Third, task-activated parcels were highly connected during rest, being significantly more connected to one another than to other regions. Fourth, resting-state connections could accurately predict illusory/fragmented task activation differences via ActFlow, which implies that these same connections were employed for shape completion. Fifth, significant dorsal attention regions could model task activity within the secondary visual network and across all remaining networks, indicating that this network may coordinate activity across networks during shape completion. Finally, primary visual cortex was involved in shape completion, but its influence was weaker and could only be inferred at a vertex-wise spatial resolution. Below, we discuss these findings in more detail, provide a sketch of how these regions might interact during shape completion, identify potential limitations, and suggest future directions.

### 4.1. A central role for visual networks in shape completion

Our results confirm past work showing the centrality of visual cortex for shape completion. The secondary visual network contained 32 percent of all significantly activated parcels and the parcel-wise task activation differences in this network—but not others—could classify task condition. *A priori* regions of interest—V4, LO1, LO2, LO3, V3CD—were all significant in at least one hemisphere on the task activation analysis; V4 and LO2 were each significant in at least one hemisphere on the vertex-wise MVPA analysis. The secondary visual region—parcel PH, discussed further below—was a standout in its consistency across subjects. Significant visual parcels were categorically more active in the illusory than fragmented condition without regard to accuracy and were spatially contiguous on the lateral surface (see swath of purple in the lateral views of Fig. 4A), suggesting that shape completion could potentially be boosted by transcranially stimulating this network. Such interventions could potentially treat conditions that impair shape completion such as developmental agnosia (deactivated mid-level visual areas), schizophrenia, brain injury (infarct/hemorrhage), or recent recovery from congenital blindness (cataract removal) (Gilaie-Dotan et al., 2009; Keane et al., 2019; Ostrovsky et al., 2009; Vuilleumier et al., 2001).

The primary visual network region, V1, was significant only on a vertex-wise MVPA analysis. A likely reason is that—according to a population receptive field mapping approach (Kok and de Lange, 2014)—the illusory shape surface region (corresponding to a portion of V1 vertices) is more activated in V1 relative to baseline and the inducer (pac-man) regions are *less* activated. Therefore, averaging across these two retinotopic region types will reveal no changes in overall activity. Our null parcel-wise and significant vertex-wise results provide some support for this view. This may also explain why, historically, the methods with the highest spatial resolution were those that provided the most convincing evidence for illusory contour formation in V1 and V2 (Grosof et al., 1993; Kok and de Lange, 2014; e.g., Lee and Nguyen, 2001) and why many lower-resolution neuroimaging studies have often failed to find effects at this level (Seghier and Vuilleumier, 2006). For example, a study using the fat/thin discrimination task with 3 mm voxels found no modulation of early visual areas (Stanley and Rubin, 2003) whereas a behaviorally similar study using 2 mm voxels and a surface-based analysis revealed effects (Maertens et al., 2008). Therefore, higher resolution fMRI may be needed to better bring out effects more fully within V1 and V2.

### 4.2. Area PH: A potential “classical region” for shape completion and a link to reading

Area PH is a recently re-defined region in the posterior temporal cortex, corresponding to the superior part of PH in the von Economo and Koskinas atlas (Glasser et al., 2016; Triarhou, 2007); it is not commonly reported in the neuroimaging literature and was not of a priori interest. Nevertheless, in our study it was the most consistently active parcel across subjects and the most frequently significant parcel within subject. Among modulated task regions, it was also the most densely connected visual parcel, communicating directly with lateral occipital cortex (V3CD), consistent with past research (Glasser et al., 2016). In light of these results, PH should be considered a candidate “classical region” for shape completion along with other more recognized areas such as lateral occipital cortex. Strong activation of PH could also explain why the fusiform face area has been occasionally reported in past studies of shape completion (Halgren et al., 2003; Larsson et al., 1999) since PH is immediately bordering the fusiform face complex and since signal leakage or improper delineation of PH would inevitably result in false positives. Finally, PH has been considered by some to be the best atlas-based alternative to the functionally-defined visual word form area (VWFA; Weiss et al., 2019). The VWFA has been shown to have high functional connectivity to the dorsal attention network (Vogel et al., 2012). Consistent with this finding, we showed that area PH was significantly connected to task-modulated dorsal attention regions in each hemisphere (MIP, LIPd, PFt, AIP, IP0); Fig. 5C). An interesting possibility is that visual shape completion may be compromised in those with dyslexia (Monzalvo et al., 2012) and that the two abilities may be related behaviorally in non-clinical populations. A related possibility is that area PH may help explain why people with schizophrenia exhibit both reading (Revheim et al., 2014) and completion deficits (Keane et al., 2019) and why patients with this psychiatric disorder are more susceptible to developmental dyslexia (Whitford et al., 2018).

### 4.3. Frontoparietal feedback to mid-level vision via the dorsal attention network

Frontoparietal network regions were differentially active in orbitofrontal, dorsolateral prefrontal, and posterior parietal cortex. Despite receiving little regard in the literature (M. M.Murray and Herrmann, 2013; Seghier and Vuilleumier, 2006), frontoparietal involvement is not wildly unexpected. In the aforementioned MEG study, peak orbitofrontal modulation from passively-viewed Kanizsa shapes arose 340 ms post stimulus onset (Halgren et al., 2003). In eight month-(but not six month-) old infants, gamma band oscillations (40 HZ) from Kanizsa shapes were generated over frontal electrodes between 240-320 ms (Csibra et al., 2000).

Frontoparietal regions may create expectation-based predictions (Bar, 2003) for amplifying less salient illusory contours and thereby improving task performance. For example, blurry lightness-induced surfaces (so-called “salient regions; Stanley and Rubin, 2003) generate a delayed LOC activation relative to standard Kanizsa shapes (Shpaner et al., 2009), potentially reflecting the brain’s late-arriving best guesses about the precise shape of the incoming stimulus. In a fat/thin discrimination behavioral study, biasing observers to see edge elements as disconnected worsened the discrimination of illusory but not fragmented shapes (Keane et al., 2012), suggesting again that noticing and using illusory contours for shape discrimination requires appropriately conceptualizing the stimulus. Top-down signals may additionally allow observers to cognitively infer (or “abstract”) missing contours that cannot be formed via illusory contour formation such as when edge elements are extremely sparse, misaligned, or misoriented (Keane, 2018; Wyatte et al., 2014). Finally, frontoparietal network may communicate with mid-level visual structures primarily by way of the dorsal attention network (Cavada and Goldman-Rakic, 1989). As evidence, all nine modulated frontoparietal parcels in our resting-state analysis were significantly connected to at least one dorsal attention region, most typically in the posterior parietal cortex.

Note that high-level feedback of the type described is compatible with a fast, automatic and overall modular illusory contour formation process (Keane, 2018). Illusory contours begin forming at 70 ms post-stimulus onset in V2 (Lee and Nguyen, 2001) and 90 ms in LOC (M. Murray et al., 2002), which is well before the arrival of higher-order feedback. Higher-order cortical feedback may be ineffectual even after its arrival, if it must compete with persistently salient bottom-up signals (Desimone and Duncan, 1995; Keane, 2018; McMains and Kastner, 2010). Parietal neglect patients with damage to inferior parietal cortex can form illusory contours (Vuilleumier, Valenza, & Landis, 2001) and people with prefrontal cortical lesions can integrate disconnected contour elements (Ciaramelli et al., 2007), suggesting again that these areas may not be necessary for forming illusory contours. Thus, frontoparietal signals—and their dorsal attention conduits—may primarily be important for performing computations on contours already formed in mid-level vision.

### 4.4. Objections, limitations, and opportunities for future research

An objection is that the illusory and fragmented conditions required observers to judge different aspects of the stimulus (orientation or shape), and so differences in “task set” rather than shape completion may explain our results. To address this objection, we first note that *any* adequate control condition will lack shape completion and require seeing the stimulus as categorically different. Therefore, it is not possible to perfectly control for task set without obliterating the difference of interest. Second, differences in task set—at least in our study— did not make our two conditions incommensurable since the two were highly correlated in accuracy and reaction time (*r*s>.7.s, *p*s<.001). These correlations are noteworthy because most visual tasks are only weakly correlated despite having high test-retest reliability (Grzeczkowski et al., 2017). Our high correlations suggest that the left/right task successfully controlled for processes that were not of central interest (visual short term memory, spatial attention, vigilance, etc.).

Another objection is that eye movement differences could have confounded our results. This objection is weakened by five considerations: 1) all subjects were repeatedly asked to fixate within and outside of the scanner; 2) pac-men locations were equidistant from fixation, equally informative within a trial, and matched between conditions, reducing the chance of systematic differences; 3) the illusory and fragmented conditions were correlated in RT and accuracy, even for the accuracy matched sub-sample (n=10; *r*s>.8, *p*s<.01), suggesting again that any possible eye movement differences impacted performance minimally; 4) saccading after stimulus onset would offer little benefit since saccade latency is ~200 ms (Sumner, 2011) and the stimuli appeared for only 250 ms at unpredictable times during a block; and, 5) there is little evidence that eye movements impact visual shape completion in non-translating displays and some evidence that it has no effect (Cox et al., 2013). Thus, while we cannot completely rule out eye movement confounds, they are unlikely to explain our results.

Past psychophysical studies have shown similar illusory and fragmented task performance (Keane et al., 2014) but in the present study the fragmented task was about 7 percent better. Why? A possible reason is that past studies required a verbal response on each trial while ours required a button press. The congruence between the left and right rotations and left and right button press may have conferred a small but consistent benefit perhaps by diminishing the likelihood of misremembering the mapping between keypress and response. We do not view this as problematic, since the effects arose when the congruence benefit was behaviorally eliminated—both in RT and accuracy. Therefore, while large task accuracy differences clearly alter the neural data (as in the easy versus hard contrast), smaller differences appear to have little effect.

Limitations are worth noting. Even though our shape completion study had certain methodological advantages (e.g., multiband, surface-based analysis), a larger sample and a higher field magnet will likely reveal additional regions, connections, and networks. A larger sample could also allow us to better ascertain correlations between parcel-wise task modulations and condition-wise performance differences. As has already been noted, the slow hemodynamic response prevents a full description of the temporal dynamics. Additional control conditions or eye movement analyses could further support the conclusions argued above.

To summarize, the present research identified a restricted set of densely-interconnected regions that were responsive to visually completed shapes. The secondary visual network— especially area PH—played a dominant role in the process, but portions of at least four other networks were also involved, suggesting that shape completion is a distributed process. The dorsal attention network parcels appeared to coordinate activity in the secondary visual network and across cortex during visual shape completion. A logical next step will be to apply neurostimulation to probe parcel-wise causal interactions or electrophysiology to assess their activity flow dynamics.

## Acknowledgments

We thank Laura Crespo, Lisa Cruz, Dillon Smith, and Megan Serody for help in recruiting participants and collecting and organizing study data. We are also indebted to Michael Harms for assistance in finalizing the pulse sequence, and Takuya Ito and Carrisa Cocuzza for providing sample code. The authors additionally acknowledge the Office of Advanced Research Computing (OARC) at Rutgers University for providing access to the Amarel cluster and associated research computing resources (http://oarc.rutgers.edu).

## Supplementary Material

### Controlling for task accuracy differences by examining a participant subse

One way that we controlled for between-task accuracy differences was simply by examining half of the participants (n=10) who had the largest (or least negative) illusory/fragmented difference. None of the results changed if we were to use 11 participants (instead of 10). If only 9 participants were used, all results were again the same, except that left LO1 and left V3CD became non-significant probably from insufficient power.

### Supplementary Figure

**Fig. S1.**
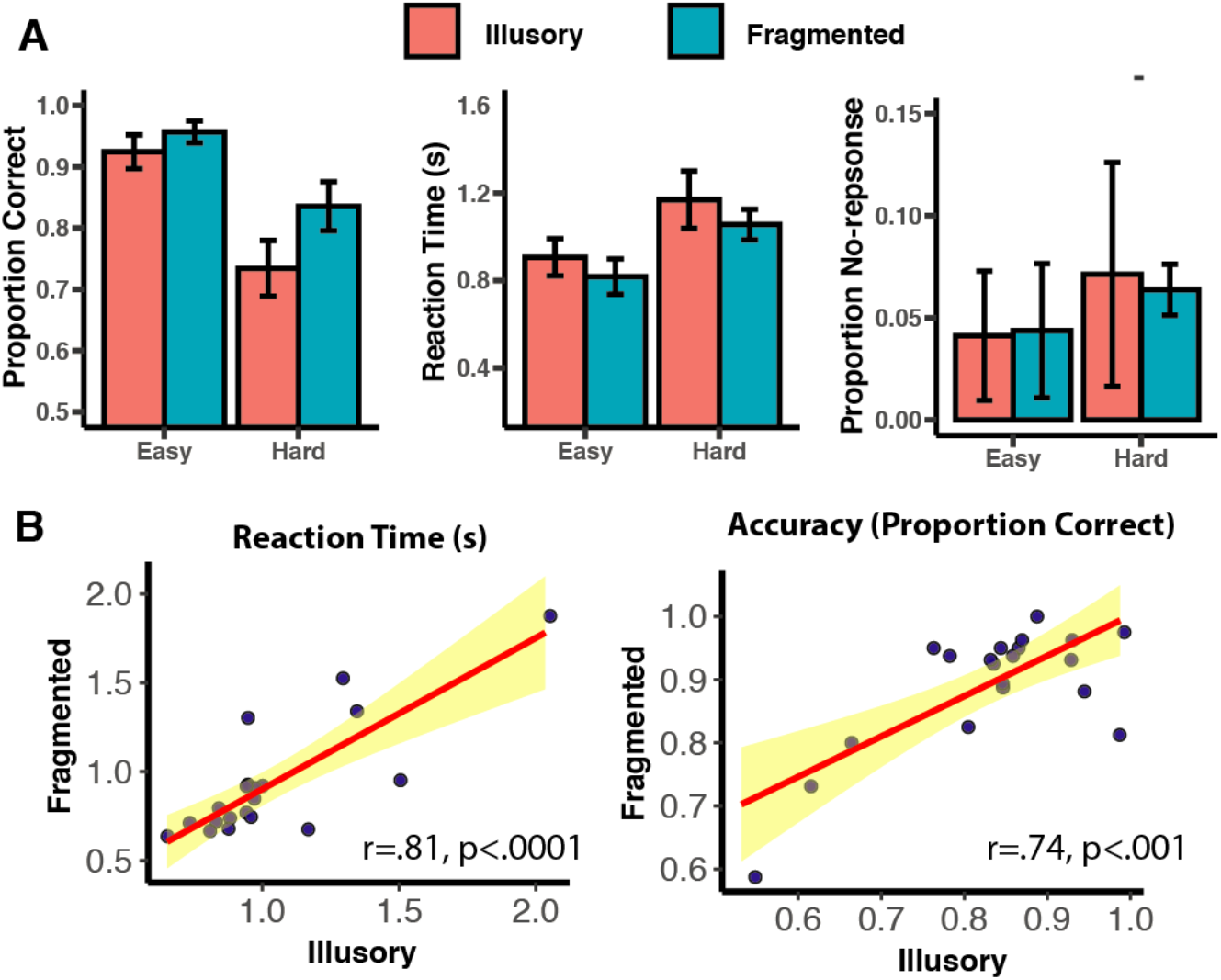
(A) Performance metrics for all conditions. (B) The two tasks were correlated in response time and accuracy.

## Footnotes

1 Shape completion effects in IT (Huxlin et al., 2000; Sáry et al., 2008) also count as evidence for classical regions since this structure is a plausible LO homologue (Orban et al., 2004).

2 The terms were “illusory contours” OR “illusory contour” OR “modal completion” OR “subjective contours” OR “subjective contour” OR “contour completion” OR “perceptual completion” OR “visual completion” OR “contour interpolation” OR “Kanizsa”; search date = 10/11/20

## References

Anticevic, A., Cole, M.W., Murray, J.D., Corlett, P.R., Wang, X.-J., Krystal, J.H., 2012. The role of default network deactivation in cognition and disease. Trends Cogn Sci (Regul Ed) 16, 584–592. doi:10.1016/j.tics.2012.10.008

Bar, M., 2003. A cortical mechanism for triggering top-down facilitation in visual object recognition. J Cogn Neurosci 15, 600–609. doi:10.1162/089892903321662976

Beck, J., 1966. Effect of orientation and of shape similarity on perceptual grouping. Perception & Psychophysics 1, 300–302.

Benjamini, Y., Hochberg, J., 1995. Controlling the false discovery rate: A practical and powerful approach to multiple testing. Journal of the Royal Statistical Society Series B-Methodological 57, 289–300.

Cavada, C., Goldman-Rakic, P.S., 1989. Posterior parietal cortex in rhesus monkey: II. Evidence for segregated corticocortical networks linking sensory and limbic areas with the frontal lobe. J. Comp. Neurol. 287, 422–445. doi:10.1002/cne.902870403

Ciaramelli, E., Leo, F., Del Viva, M.M., Burr, D.C., Ladavas, E., 2007. The contribution of prefrontal cortex to global perception. Exp Brain Res 181, 427–434. doi:10.1007/s00221-007-0939-7

Ciric, R., Wolf, D.H., Power, J.D., Roalf, D.R., Baum, G.L., Ruparel, K., Shinohara, R.T., Elliott, M.A., Eickhoff, S.B., Davatzikos, C., Gur, R.C., Gur, R.E., Bassett, D.S., Satterthwaite, T.D., 2017. Benchmarking of participant-level confound regression strategies for the control of motion artifact in studies of functional connectivity. Neuroimage 154, 174–187. doi:10.1016/j.neuroimage.2017.03.020

Cole, M.W., Bassett, D.S., Power, J.D., Braver, T.S., Petersen, S.E., 2014. Intrinsic and task-evoked network architectures of the human brain. Neuron 83, 238–251. doi:10.1016/j.neuron.2014.05.014

Cole, M.W., Ito, T., Bassett, D.S., Schultz, D.H., 2016. Activity flow over resting-state networks shapes cognitive task activations. Nature Neuroscience 19, 1718–1726. doi:10.1038/nn.4406

Cox, M.A., Schmid, M.C., Peters, A.J., Saunders, R.C., Leopold, D.A., Maier, A., 2013. Receptive field focus of visual area V4 neurons determines responses to illusory surfaces. Proc Natl Acad Sci USA 110, 17095–17100. doi:10.1073/pnas.1310806110/-/DCSupplemental

Csibra, G., Davis, G., Spratling, M.W., Johnson, M.H., 2000. Gamma oscillations and object processing in the infant brain. Science 290, 1582–1585.

Desimone, R., Duncan, J., 1995. Neural mechanisms of selective visual attention. Annu Rev Neurosci 18, 193–222.

Doniger, G.M., Foxe, J.J., Murray, M.M., Higgins, B.A., Javitt, D.C., 2002. Impaired visual object recognition and dorsal/ventral stream interaction in schizophrenia. Arch Gen Psychiatry 59, 1011–1020.

Esteban, O., Birman, D., Schaer, M., Koyejo, O.O., Poldrack, R.A., Gorgolewski, K.J., 2017. MRIQC: Advancing the automatic prediction of image quality in MRI from unseen sites. PLOS ONE 12, e0184661. doi:10.1371/journal.pone.0184661.s001

Foxe, J.J., Murray, M.M., Javitt, D.C., 2005. Filling-in in schizophrenia: a high-density electrical mapping and source-analysis investigation of illusory contour processing. Cereb Cortex 15, 1914–1927. doi:10.1093/cercor/bhi069

Gilaie-Dotan, S., Perry, A., Bonneh, Y., Malach, R., Bentin, S., 2009. Seeing with profoundly deactivated mid-level visual areas: non-hierarchical functioning in the human visual cortex. Cerebral Cortex 19, 1687–1703. doi:10.1093/cercor/bhn205

Glasser, M.F., Coalson, T.S., Robinson, E.C., Hacker, C.D., Harwell, J., Yacoub, E., Uğurbil, K., Andersson, J., Beckmann, C.F., Jenkinson, M., Smith, S.M., Van Essen, D.C., 2016. A multi-modal parcellation of human cerebral cortex. Nature 536, 171–178. doi:10.1038/nature18933

Glasser, M.F., Sotiropoulos, S.N., Wilson, J.A., Coalson, T.S., Fischl, B., Andersson, J.L., Xu, J., Jbabdi, S., Webster, M., Polimeni, J.R., Van Essen, D.C., Jenkinson, M., Consortium, F.T.W.-M.H., 2013. The minimal preprocessing pipelines for the Human Connectome Project. Neuroimage 80, 105–124. doi:10.1016/j.neuroimage.2013.04.127

Gold, J.M., Murray, R.F., Bennett, P.J., Sekuler, A.B., 2000. Deriving behavioural receptive fields for visually completed contours. Current Biology 10, 663–666.

Grosof, D.H., Shapley, R.M., Hawken, M.J., 1993. Macaque V1 neurons can signal “illusory” contours. Nature 365, 550–552. doi:10.1038/365550a0

Grzeczkowski, L., Clarke, A.M., Francis, G., Mast, F.W., Herzog, M.H., 2017. About individual differences in vision. Vision Research 141, 282–292. doi:10.1016/j.visres.2016.10.006

Halgren, E., Mendola, J., Chong, C.D.R., Dale, A.M., 2003. Cortical activation to illusory shapes as measured with magnetoencephalography. Neuroimage 18, 1001–1009.

Haynes, J.-D., 2015. A Primer on Pattern-Based Approaches to fMRI: Principles, Pitfalls, and Perspectives. Neuron 87, 257–270. doi:10.1016/j.neuron.2015.05.025

Huxlin, K.R., Saunders, R.C., Marchionini, D., Pham, H.A., Merigan, W.H., 2000. Perceptual deficits after lesions of inferotemporal cortex in macaques. Cereb Cortex 10, 671–683. doi:10.1093/cercor/10.7.671

Iacaruso, M.F., Gasler, I.T., Hofer, S.B., 2017. Synaptic organization of visual space in primary visual cortex. Nature 547, 449–452. doi:10.1038/nature23019

Ito, T., Kulkarni, K.R., Schultz, D.H., Mill, R.D., Chen, R.H., Solomyak, L.I., Cole, M.W., 2017. Cognitive task information is transferred between brain regions via resting-state network topology. Nat Comms 8, 1027. doi:10.1038/s41467-017-01000-w

Ji, J.L., Spronk, M., Kulkarni, K., Repovs, G., Anticevic, A., Cole, M.W., 2019. Mapping the human brain’s cortical-subcortical functional network organization. Neuroimage 185, 35–57. doi:10.1016/j.neuroimage.2018.10.006

Keane, B.P., 2018. Contour interpolation: A case study in Modularity of Mind. Cognition 174, 1–18. doi:10.1016/j.cognition.2018.01.008

Keane, B.P., Joseph, J., Silverstein, S.M., 2014. Late, not early, stages of Kanizsa shape perception are compromised in schizophrenia. Neuropsychologia 56, 302–311. doi:10.1016/j.neuropsychologia.2014.02.001

Keane, B.P., Lu, H., Kellman, P.J., 2007. Classification images reveal spatiotemporal contour interpolation. Vision Research 47, 3460–3475. doi:10.1016/j.visres.2007.10.003

Keane, B.P., Lu, H., Papathomas, T.V., Silverstein, S.M., Kellman, P.J., 2012. Is interpolation cognitively encapsulated? Measuring the effects of belief on Kanizsa shape discrimination and illusory contour formation. Cognition 123, 404–418. doi:10.1016/j.cognition.2012.02.004

Keane, B.P., Paterno, D., Kastner, S., Krekelberg, B., Silverstein, S.M., 2019. Intact illusory contour formation but equivalently impaired visual shape completion in first- and later-episode schizophrenia. J Abnorm Psychol 128, 57–68. doi:10.1037/abn0000384

Kellman, P.J., Shipley, T., 1991. A theory of visual interpolation in object perception. Cogn Psychol 23, 141.

Kok, P., de Lange, F.P., 2014. Shape Perception Simultaneously Up- and Downregulates Neural Activity in the Primary Visual Cortex. Current Biology 24, 1531–1535. doi:10.1016/j.cub.2014.05.042

Kruggel, F., Herrmann, C.S., Wiggins, C.J., Cramon von, D.Y., 2001. Hemodynamic and Electroencephalographic Responses to Illusory Figures: Recording of the Evoked Potentials during Functional MRI. Neuroimage 14, 1327–1336. doi:10.1006/nimg.2001.0948

Larsson, J., Amunts, K., Gulyás, B., Malikovic, A., Zilles, K., Roland, P.E., 1999. Neuronal correlates of real and illusory contour perception: functional anatomy with PET. Eur J Neurosci 11, 4024–4036. doi:10.1046/j.1460-9568.1999.00805.x

Lee, T., Nguyen, M., 2001. Dynamics of subjective contour formation in the early visual cortex. Proceedings of the National Academy of Sciences 98, 1907–1911. doi:10.1073/pnas.031579998

Maertens, M., Pollmann, S., Hanke, M., Mildner, T., Möller, H., 2008. Retinotopic activation in response to subjective contours in primary visual cortex. Front. Hum. Neurosci. 2, 2. doi:10.3389/neuro.09.002.2008

Malikovic, A., Amunts, K., Schleicher, A., Mohlberg, H., Kujovic, M., Palomero-Gallagher, N., Eickhoff, S.B., Zilles, K., 2016. Cytoarchitecture of the human lateral occipital cortex: mapping of two extrastriate areas hOc4la and hOc4lp. Brain Struct Funct 221, 1877–1897. doi:10.1007/s00429-015-1009-8

McMains, S.A., Kastner, S., 2010. Defining the units of competition: influences of perceptual organization on competitive interactions in human visual cortex. J Cogn Neurosci 22, 2417–2426. doi:10.1162/jocn.2009.21391

Mendola, J.D., Dale, A.M., Fischl, B., Liu, A.K., Tootell, R.B., 1999. The representation of illusory and real contours in human cortical visual areas revealed by functional magnetic resonance imaging. J Neurosci 19, 8560–8572.

Merboldt, K.-D., Finsterbusch, J., Frahm, J., 2000. Reducing Inhomogeneity Artifacts in Functional MRI of Human Brain Activation—Thin Sections vs Gradient Compensation. Journal of Magnetic Resonance 145, 184–191. doi:10.1006/jmre.2000.2105

Mill, R.D., Gordon, B.A., Balota, D.A., Cole, M.W., 2020. Predicting dysfunctional age-related task activations from resting-state network alterations. Neuroimage 117167–37. doi:10.1016/j.neuroimage.2020.117167

Monzalvo, K., Fluss, J., Billard, C., Dehaene, S., Dehaene-Lambertz, G., 2012. Cortical networks for vision and language in dyslexic and normal children of variable socio-economic status. Neuroimage 61, 258–274. doi:10.1016/j.neuroimage.2012.02.035

Mur, M., Bandettini, P.A., Kriegeskorte, N., 2009. Revealing representational content with pattern-information fMRI--an introductory guide. Soc Cogn Affect Neurosci 4, 101–109. doi:10.1093/scan/nsn044

Murray, M., Wylie, G., Higgins, B., Javitt, D., Schroeder, C., Foxe, J., 2002. The spatiotemporal dynamics of illusory contour processing: combined high-density electrical mapping, source analysis, and functional magnetic resonance imaging. Journal of Neuroscience 22, 5055.

Murray, M.M., Foxe, D.M., Javitt, D.C., Foxe, J.J., 2004. Setting boundaries: brain dynamics of modal and amodal illusory shape completion in humans. Journal of Neuroscience 24, 6898–6903. doi:10.1523/JNEUROSCI.1996-04.2004

Murray, M.M., Herrmann, C.S., 2013. Illusory contours: a window onto the neurophysiology of constructing perception. Trends Cogn Sci (Regul Ed). doi:10.1016/j.tics.2013.07.004

Murray, M.M., Imber, M.L., Javitt, D.C., Foxe, J.J., 2006. Boundary Completion Is Automatic and Dissociable from Shape Discrimination. Journal of Neuroscience 26, 12043–12054. doi:10.1523/JNEUROSCI.3225-06.2006

Nieder, A., 2002. Seeing more than meets the eye: processing of illusory contours in animals. J Comp Physiol A Neuroethol Sens Neural Behav Physiol 188, 249–260. doi:10.1007/s00359-002-0306-x

Orban, G.A., Van Essen, D., Vanduffel, W., 2004. Comparative mapping of higher visual areas in monkeys and humans. Trends Cogn Sci (Regul Ed) 8, 315–324. doi:10.1016/j.tics.2004.05.009

Ostrovsky, Y., Meyers, E., Ganesh, S., Mathur, U., Sinha, P., 2009. Visual parsing after recovery from blindness. Psychol Sci 20, 1484–1491. doi:10.1111/j.1467-9280.2009.02471.x

Peirce, J.W., 2007. PsychoPy—Psychophysics software in Python. J Neurosci Methods 162, 8–13. doi:10.1016/j.jneumeth.2006.11.017

Pelli, D.G., 1997. The VideoToolbox software for visual psychophysics: transforming numbers into movies. Spat Vis 10, 437–442.

Pillow, J., Rubin, N., 2002. Perceptual completion across the vertical meridian and the role of early visual cortex. Neuron 33, 805–813.

Power, J.D., Barnes, K.A., Snyder, A.Z., Schlaggar, B.L., Petersen, S.E., 2012. Spurious but systematic correlations in functional connectivity MRI networks arise from subject motion. Neuroimage 59, 2142–2154. doi:10.1016/j.neuroimage.2011.10.018

Power, J.D., Cohen, A.L., Nelson, S.M., Wig, G.S., Barnes, K.A., Church, J.A., Vogel, A.C., Laumann, T.O., Miezin, F.M., Schlaggar, B.L., Petersen, S.E., 2011. Functional Network Organization of the Human Brain. Neuron 72, 665–678. doi:10.1016/j.neuron.2011.09.006

Revheim, N., Corcoran, C.M., Dias, E., Hellmann, E., Martinez, A., Butler, P.D., Lehrfeld, J.M., DiCostanzo, J., Albert, J., Javitt, D.C., 2014. Reading deficits in schizophrenia and individuals at high clinical risk: relationship to sensory function, course of illness, and psychosocial outcome. Am J Psychiatry 171, 949–959. doi:10.1176/appi.ajp.2014.13091196

Ringach, D., Shapley, R., 1996. Spatial and temporal properties of illusory contours and amodal boundary completion. Vision Research 36, 3037–3050.

Sáry, G., Köteles, K., Kaposvári, P., Lenti, L., Csifcsák, G., Frankó, E., Benedek, G., Tompa, T., 2008. The representation of Kanizsa illusory contours in the monkey inferior temporal cortex. European Journal of Neuroscience 28, 2137–2146. doi:10.3758/BF03198792

Schultz, D.H., Ito, T., Solomyak, L.I., Chen, R.H., Mill, R.D., Anticevic, A., Cole, M.W., 2018. Global connectivity of the fronto-parietal cognitive control network is related to depression symptoms in the general population. Netw Neurosci 3, 107–123. doi:10.1162/netn_a_00056

Seghier, M.L., Vuilleumier, P., 2006. Functional neuroimaging findings on the human perception of illusory contours. Neurosci Biobehav Rev 30, 595–612. doi:10.1016/j.neubiorev.2005.11.002

Shpaner, M., Murray, M.M., Foxe, J.J., 2009. Early processing in the human lateral occipital complex is highly responsive to illusory contours but not to salient regions. European Journal of Neuroscience 30, 2018–2028. doi:10.1111/j.1460-9568.2009.06981.x

Smith, S.M., Beckmann, C.F., Andersson, J., Auerbach, E.J., Bijsterbosch, J., Douaud, G., Duff, E., Feinberg, D.A., Griffanti, L., Harms, M.P., Kelly, M., Laumann, T., Miller, K.L., Moeller, S., Petersen, S., Power, J., Salimi-Khorshidi, G., Snyder, A.Z., Vu, A.T., Woolrich, M.W., Xu, J., Yacoub, E., Uğurbil, K., Van Essen, D.C., Glasser, M.F., 2013. Resting-state fMRI in the Human Connectome Project. Neuroimage 80, 144–168. doi:10.1016/j.neuroimage.2013.05.039

Spronk, M., Kulkarni, K., Ji, J.L., Keane, B.P., Anticevic, A., Cole, M.W., 2018. A whole-brain and cross-diagnostic perspective on functional brain network dysfunction. BioRxiv 1–23. doi:10.1101/326728

Stanley, D.A., Rubin, N., 2003. fMRI activation in response to illusory contours and salient regions in the human lateral occipital complex. Neuron 37, 323–331.

Stark, D.E., Margulies, D.S., Shehzad, Z.E., Reiss, P., Kelly, A.M.C., Uddin, L.Q., Gee, D.G., Roy, A.K., Banich, M.T., Castellanos, F.X., Milham, M.P., 2008. Regional variation in interhemispheric coordination of intrinsic hemodynamic fluctuations. Journal of Neuroscience 28, 13754–13764. doi:10.1523/JNEUROSCI.4544-08.2008

Sumner, P., 2011. Determinants of saccade latency, in: Liversedge, S., Gilchrist, I., Everling, S. (Eds.), The Oxford Handbook of Eye Movements. books.google.com, New York, pp. 411–424.

Triarhou, L.C., 2007. The Economo-Koskinas atlas revisited: Cytoarchitectonics and functional context. Stereotact Funct Neurosurg 85, 195–203. doi:10.1159/000103258

Valenza, E., Bulf, H., 2010. Early development of object unity: evidence for perceptual completion in newborns. Developmental Sci 1–10. doi:10.1111/j.1467-7687.2010.01026.x

Vogel, A.C., Miezin, F.M., Petersen, S.E., Schlaggar, B.L., 2012. The putative visual word form area is functionally connected to the dorsal attention network. Cerebral Cortex 22, 537–549. doi:10.1093/cercor/bhr100

Vuilleumier, P., Valenza, N., Landis, T., 2001. Explicit and implicit perception of illusory contours in unilateral spatial neglect: behavioural and anatomical correlates of preattentive grouping mechanisms. Neuropsychologia 39, 597–610.

Weiss, F., Greenlee, M.W., Volbert, G., 2019. No atypical white-matter structures in grapheme- or color-sensitive areas in synesthetes. BioRxiv. doi:10.1101/618611

Whitford, V., O’Driscoll, G.A., Titone, D., 2018. Reading deficits in schizophrenia and their relationship to developmental dyslexia: A review. Schizophr Res 193, 11–22. doi:10.1016/j.schres.2017.06.049

Wokke, M.E., Vandenbroucke, A.R.E., Scholte, H.S., Lamme, V.A.F., 2013. Confuse your illusion: feedback to early visual cortex contributes to perceptual completion. Psychological Science 24, 63–71. doi:10.1177/0956797612449175

Wyatte, D., Jilk, D.J., O’Reilly, R.C., 2014. Early recurrent feedback facilitates visual object recognition under challenging conditions. Front. Psychol. 5, 760–10. doi:10.3389/fpsyg.2014.00674

